# Chromatin Structures from Integrated AI and Polymer Physics Model

**DOI:** 10.1101/2024.11.27.624905

**Authors:** Eric R Schultz, Soren Kyhl, Rebecca Willett, Juan J de Pablo

## Abstract

The physical organization of the genome in three-dimensional space regulates many biological processes, including gene expression and cell differentiation. Three-dimensional characterization of genome structure is critical to understanding these biological processes. Direct experimental measurements of genome structure are challenging; computational models of chromatin structure are therefore necessary. We develop an approach that combines a particle-based chromatin polymer model, molecular simulation, and machine learning to efficiently and accurately estimate chromatin structure from *indirect* measures of genome structure. More specifically, we introduce a new approach where the interaction parameters of the polymer model are extracted from experimental Hi-C data using a graph neural network (GNN). We train the GNN on simulated data from the underlying polymer model, avoiding the need for large quantities of experimental data. The resulting approach accurately estimates chromatin structures across all chromosomes and across several experimental cell lines despite being trained almost exclusively on simulated data. The proposed approach can be viewed as a general framework for combining physical modeling with machine learning, and it could be extended to integrate additional biological data modalities. Ultimately, we achieve accurate and high-throughput estimations of chromatin structure from Hi-C data, which will be necessary as experimental methodologies, such as single-cell Hi-C, improve.

## 1 Introduction

Advances in the ability to measure genome structure have gradually established that the three-dimensional organization of the genome is critical to the regulation of gene expression [19, 31]. Existing experimental tools for characterizing genome structure fall into two fundamental categories: microscopy-based methods and sequencing-based methods. Microscopy methods can directly measure the 3D chromatin structures of individual cells by using fluorescence in situ hybridization (FISH) tags to target specific genomic loci [4, 49]. Currently, technical challenges limit these experiments to measuring thousands of loci per experiment [49]. As a consequence, these experiments can either image a small genomic region at high resolution or a large region at low resolution, but not both.

On the other hand, sequencing technologies, such as Hi-C, can measure genome-wide chromatin organization at very high resolution, albeit indirectly [26, 43]. Hi-C quantifies three-dimensional chromatin organization by measuring the contact frequency between genomic regions. The resulting contact matrix (also referred to as a contact map) contains the population-averaged contact frequency between pairs of genomic loci. Patterns in Hi-C data provide evidence for genome structural properties such as loops [42, 43], topologically-associated domains (TADs) [10, 35], and compartmentalization [16, 26]. However, while Hi-C can measure contact frequencies at high resolution, estimating the three-dimensional structure of chromatin from Hi-C data has been a major challenge limiting the utility of Hi-C methods.

Computational models of chromatin structure provide a useful tool for elucidating the full three-dimensional structures of chromatin from Hi-C data [28, 36]. We rely on particle-based polymer models. Such models of chromatin have been shown to reproduce many properties of its structure [9, 25, 41, 57]. In these models, each particle typically represents between 5 kb and 1 Mb of DNA [36]. At this coarse of resolution, parametrization by traditional ‘bottom-up’ coarse-graining of atomic-scale or nucleosome-scale DNA models has not yet been accomplished [32, 50]. Instead, these models have been parametrized ‘top-down’ by inferring thermodynamic parameters from a target experimental contact map [36].

Polymer modeling approaches in this framework can leverage the maximum entropy principle [18] to iteratively optimize thermodynamic parameters until the simulated chromatin ensemble best reproduces the experimental contact map [9, 41, 57] (reviewed in [28]). However, the iterative nature of this optimization procedure is a major computational bottleneck in the simulation procedure. Faster methods are essential for exploring variations in chromatin structure across cell types, analyzing structural diversity in single-cell variants of Hi-C data, assessing the effect of epigenetic factors on chromatin structure, and other large-scale investigations of variations in structure. As an alternative to the maximum entropy approach, analytical methods for estimating parameters from Hi-C data are an emerging research area [45, 48]. While existing approaches are efficient, they rely on a reference homopolymer system, which limits their efficacy. Standard homopolymer potentials do not reproduce experimental chromatin contact probability scaling. As a consequence, the resulting estimated structures are less accurate than structures estimated from models which use the maximum entropy approach.

Recent advances in machine learning methods, such as graph transformers [46], which combine graph neural networks [59] with attention mechanisms [53], present an opportunity for improvement. We show that adapting these advances in machine learning tools to the task of chromatin structure prediction can dramatically accelerate structure estimation from Hi-C data. Graph neural networks are a promising architecture because the Hi-C contact map can be considered as a weighted adjacency matrix of a graph. Some existing methods that treat Hi-C data as a graph include Higashi [58], a hypergraph neural network developed for imputation of sparse single-cell Hi-C data, and Sub-Compartment Identifier (SCI) [2], which uses a graph embedding step to predict chromatin sub-compartments from Hi-C data. Transformers are a class of neural networks that rely on attention mechanisms and have been popularized by their recent success in large language models [6, 37]. Beyond language models, AlphaFold and its successor, AlphaFold 2, are graph transformers successful in the protein structure prediction problem [1, 20, 24]. We use a graph transformer with a unique approach that leverages state-of-the-art polymer models of chromatin structure.

Specifically, **we develop a graph neural network to predict the thermodynamic parameters of a polymer model from experimental Hi-C data, and these parameters allow us to simulate chromatin structures from the corresponding polymer model**. A key contribution of our approach is that we train the neural network exclusively on simulated data from the polymer model, circumventing the need for large quantities of high-quality experimental data. Further, we train our neural network to predict the interaction parameters of the polymer model rather than directly predicting a single structure. Compared to an approach that predicts only a single consensus structure, sampling structures via the polymer model better reflects both (a) uncertainty in structure estimates and (b) the dynamic nature of chromatin. Compared to using the iterative maximum entropy procedure to estimate the parameter of our polymer model, our GNN approach is 6.5x faster but yields structures of comparable quality.

## 2 Results

### 2.1 Overview of Approach

Broadly speaking, we aim to estimate the underlying chromatin structure from experimental Hi-C data [Fig. 1A]. We use a particle-based physics model where we model chromatin as a heteropolymer of *m* particles; each particle represents 50 kb of DNA [Fig. 1B]. Particles interact in our model in a pairwise fashion controlled by an interaction energy matrix, 𝒰, which contains an interaction energy, 𝒰_*ij*_, for every pair of particles *i* and *j*. To determine 𝒰, we train a graph neural network (GNN) that inputs an experimental contact map and outputs the estimated interaction parameters, 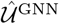, of the model [Fig. 1C]. Given the estimated interaction parameters, we use our polymer model to generate a distribution of structures. If the generated structures are accurate, we will be able to calculate a simulated contact map from these structures [Eq. (4)] that closely resembles the target experimental contact map.

**Figure 1:**
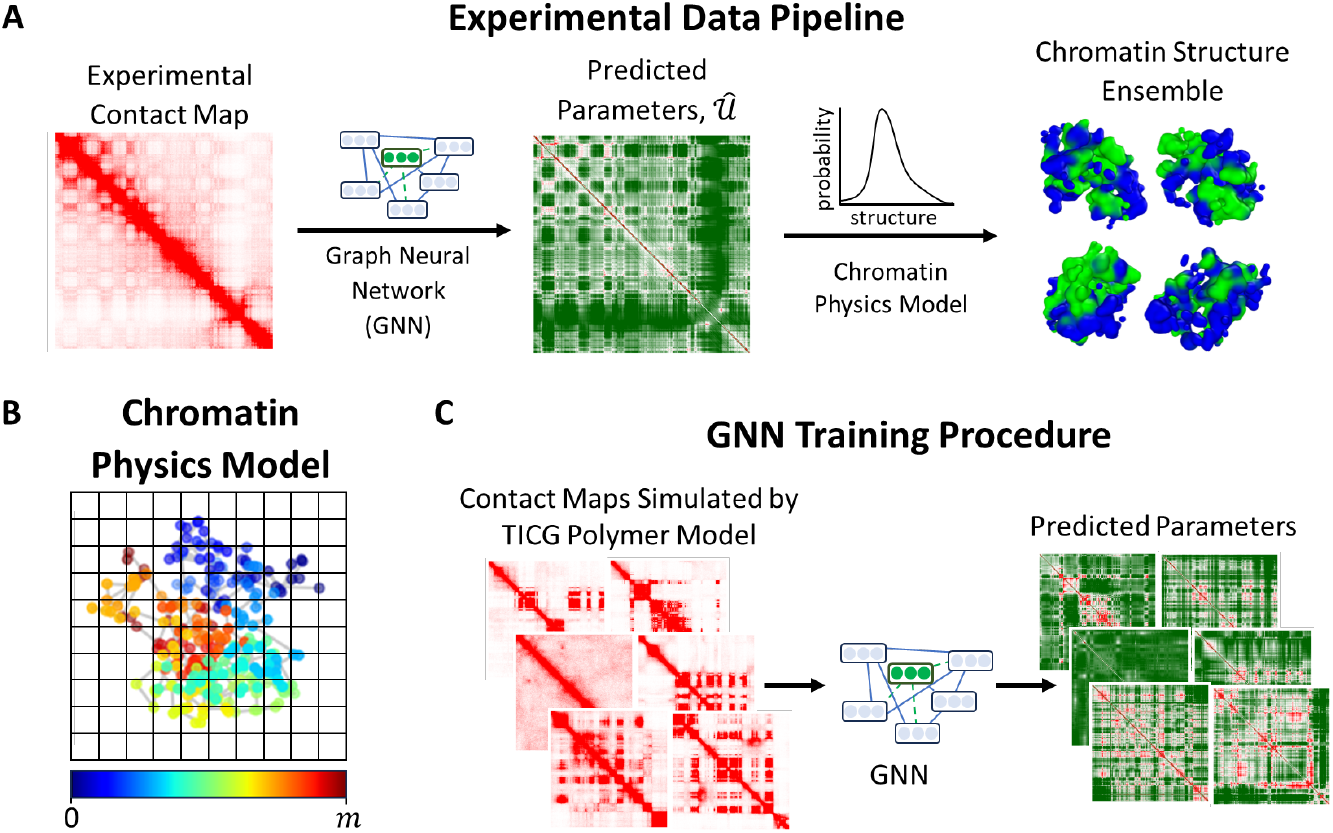
**A** Schematic overview of the experimental data pipeline. Given an experimental contact map, we construct a probability distribution over chromatin structures corresponding to our chromatin polymer model. The graph neural network (GNN) uses the contact map as an input and predicts interaction parameters, 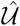, of our chromatin physics model. Given these parameters, the polymer model samples chromatin structures at random. Structures are colored by A/B compartmentalization as determined by the sign of the first principal component of the experimental genomic-distance normalized contact map. **B** Schematic overview of GNN training procedure. We train the GNN on simulated contact maps generated by the chromatin physics model where we know the ground truth interaction parameters. **C** Schematic illustration of chromatin polymer model. Particles are colored according to their position along the polymer to aid visualization. Particles in the same grid cell are defined as in contact. Grid size is not to scale.

Obtaining sufficient data to train the neural network is a major challenge, and a key contribution of our approach is that we are able to train the GNN on simulated data from our polymer model. Briefly, we first use the existing maximum entropy (ME) optimization to estimate 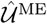 for 33 experimental contact maps all from sub-regions of the odd chromosomes from the IMR90 cell line. Then, we draw synthetic polymer model parameters from probability distributions fitted to these parameter estimates. Finally, we conduct simulations using our polymer model and the synthetic parameters to yield realistic simulated contact maps. Using this dataset, we train the GNN to predict the synthetic polymer model parameters. Intuitively, the GNN is learning to estimate the thermodynamic parameters that will yield a structural ensemble that best reproduces the experimental contact map.

After training the GNN, we can input an arbitrary 50 kb resolution experimental contact map and use the parameters, 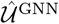, predicted by the GNN to simulate the corresponding chromatin ensemble [Fig. 1]. We will contrast this approach for simulating chromatin structures (GNN approach) with using the maximum entropy optimization to estimate the simulation parameters (maximum entropy approach). Pseudocode for both of these approaches can be found in Section S1; maximum entropy approach: Algorithm S4; GNN approach: Algorithm S5.

### 2.2 Performance on Experimental Hi-C Data

To demonstrate the accuracy and efficiency of our GNN approach, we model the chromatin structures corresponding to 40 experimental contact maps from a test set comprising the even chromosomes of the IMR90 cell line. For each experimental contact map, we use both the GNN and the maximum entropy approach to estimate the parameters of our polymer model. We compare the resulting simulated contact maps and the total simulation time of each method. We find that polymer simulations using the parameters estimated by the GNN accurately reproduce experimental contact maps despite requiring an order of magnitude less simulation time than the maximum entropy approach.

An example result for a 25.6 Mb region of GM12878 Chr2 is shown in Fig. 2. The simulated contact maps closely resemble the experimental contact map visually [Fig. 2A]. Both simulations clearly reproduce the A/B compartmentalization [26] of the experimental contact map. The A/B compartmentalization can be seen visually as the plaid pattern in the Hi-C contact map, and is captured in the first principal component of the genomic-distance normalized contact map [Eq. (5)] [Fig. 2C]. The contact probability scaling as a function of genomic distance reproduces the experimental scaling [Fig. 2B]. An example structure from the GNN simulation is shown in Fig. 2D. The structure is colored according to the A/B compartmentalization as measured by the sign of the first principal component of the genomic-distance normalized contact map. The A compartment (green) and B compartment (blue) are seen to separate in three-dimensional space. The simulated contact maps are computed from 30,000 such structures [Eq. (4)].

**Figure 2:**
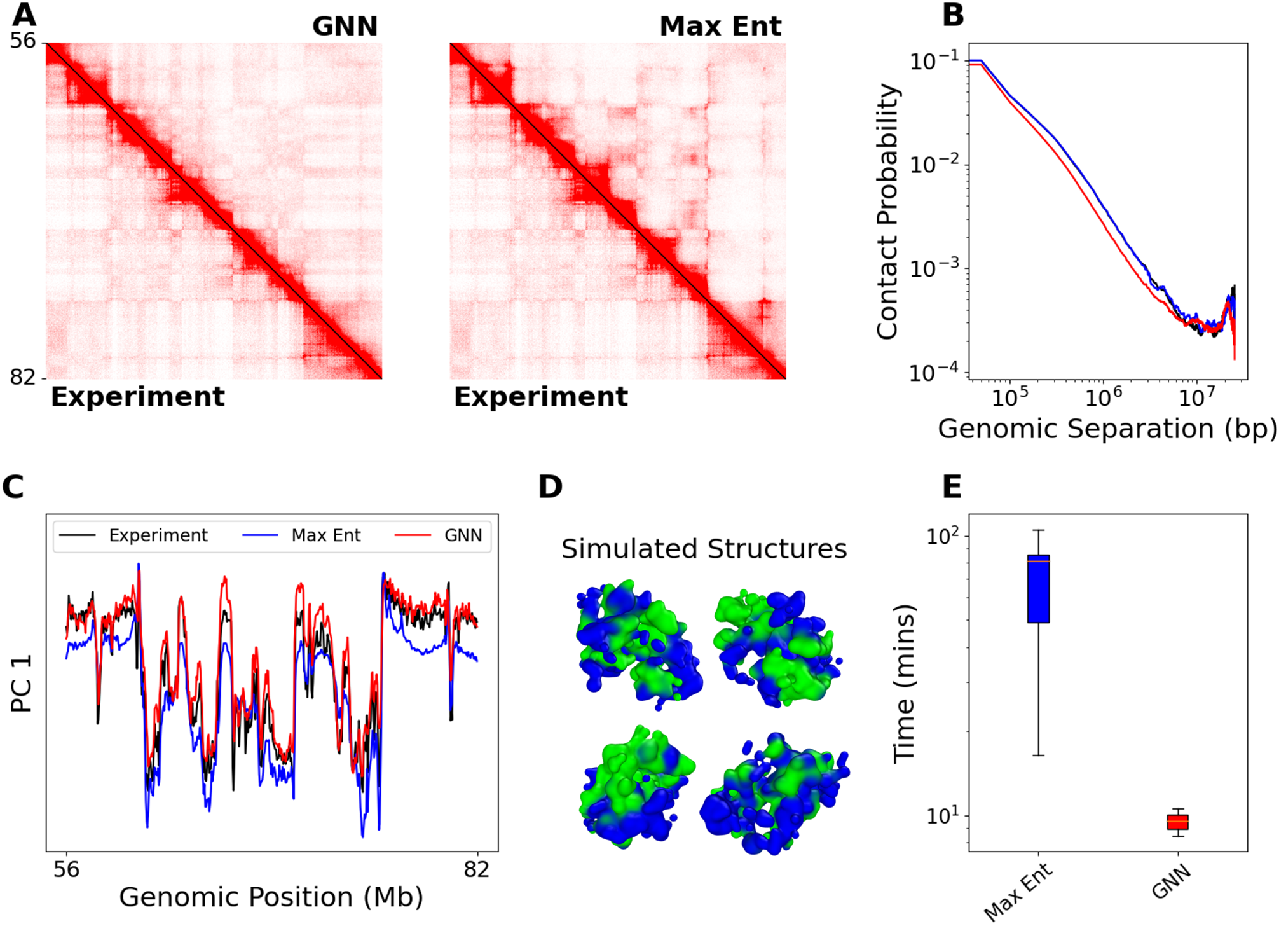
Chromatin simulation using GNN approach accurately reproduces experimental contact map. **A** Contact maps of IMR90 Chr2:56.3-81.9 Mb. The lower triangle shows the experimental contact map. The color bar is the same for both subfigures. *Left:* The upper triangle shows the GNN simulated contact map. *Right:* The upper triangle shows the maximum entropy simulated contact map. **B-C** Comparison of the experiment (black), maximum entropy simulation (blue), and GNN simulation (red). **B** Comparison of contact probability scaling as a function of genomic separation on a log-log scale. **C** Comparison of first principal component of genomic-distance normalized contact maps. **D** A representative structure from GNN simulation. Colored by A/B compartmentalization. **E** Comparison of total run time in minutes for all contact maps in the experimental test set (*n* = 40). Boxes represent the 25th and 75th quantiles, and the solid line indicates the median. Whiskers extend from the box by 1.5 times the interquartile range.

Notably, the GNN approach is 6.5 times faster than the maximum entropy approach on average [Fig. 2E]. We report the average simulation time over an experimental test set of 40 contact maps from the even chromosomes of the IMR90 cell line. The reported time only considers the time required to sample structures with polymer model [Algorithm S1] as the time of any other step is negligible in comparison. For the maximum entropy approach, the reported time includes all simulation time required for the maximum entropy optimization to converge, as well as a final simulation with the converged parameters (lines 6 and 11 in Algorithm S4). The GNN approach circumvents the maximum entropy optimization entirely. Therefore, the time reported for the GNN approach is only the simulation time using the interaction parameters estimated by the GNN (line 2 in Algorithm S5). The average time for simulations using the GNN approach is 10.3 ± 0.2 minutes. The maximum entropy method has a larger average simulation time of 66.5 ±4.0 minutes.

In Table 1, we show include quantitative results averaged over the 40 contact maps from the even chromosomes of the IMR90 cell line. Since the maximum entropy approach depends on an optimization procedure, we include two definitions of convergence in Table 1. The maximum entropy optimization is terminated based on a convergence criterion of *ε* = 10^−2^ or *ϵ* = 10^−3^ [Section S2.3.4]. Since the two convergence criteria yield comparably accurate simulated contact maps, we restrict the text to the results from *ϵ* = 10^−2^, which is faster. Note that the maximum entropy approach is not guaranteed to converge. We find that all optimizations converge for *ϵ* = 10^−2^ and 39 out of 40 converge for *ϵ* = 10^−3^. Further tuning of the user-defined parameters of the maximum entropy optimization and of the number of structures sampled via the polymer model can improve the convergence success rate.

**Table 1:** Average results for 40 experimental contact maps from even chromosomes of IMR90. The maximum entropy optimization uses a convergence criterion of either *ϵ* = 10^−2^ or *ϵ* = 10^−3^. SCC is the stratum-adjusted correlation coefficient [55]. HiC-Spector is the HiC-Spector metric from [54]. Corr PC1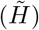 is the Pearson correlation between the first principal component of the genomic-distance normalized contact maps, 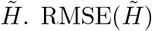 is the root-mean-squared error between the genomic-distance normalized contact maps. Simulation Time is each method’s total simulation time in minutes. See text for details [Section 2.2]. All values are mean ±standard error. *1 out of 40 maximum entropy simulations failed to meet the convergence criterion of *ϵ* = 10^−3^; we show results for the remaining 39.

| Method | SCC | HiC-Spector | Corr PC1($\tilde{H}$) | RMSE($\tilde{H}$) | Simulation Time |
| --- | --- | --- | --- | --- | --- |
| GNN | $0.72 \pm 0.01$ | $0.51 \pm 0.02$ | $0.90 \pm 0.01$ | $0.65 \pm 0.01$ | $10.3 \pm 0.2$ |
| Maximum Entropy ($\epsilon=1e-2$) | $0.81 \pm 0.01$ | $0.49 \pm 0.02$ | $0.78 \pm 0.03$ | $1.04 \pm 0.03$ | $66.5 \pm 4.0$ |
| Maximum Entropy ($\epsilon=1e-3$)* | $0.84 \pm 0.01$ | $0.51 \pm 0.02$ | $0.80 \pm 0.03$ | $0.72 \pm 0.02$ | $212.4 \pm 13.9$ |

We compare contact maps using two metrics from the Hi-C literature [54, 55]. [Table 1]. See Section S6 for further discussion of these metrics and for implementation details. The first metric is the stratum-adjusted correlation coefficient (SCC) [55]. The SCC is a weighted average of the Pearson correlations between pairs of off-diagonals of two contact maps. The simulated contact maps from simulations using the GNN approach have an SCC of 0.72 ± 0.01 (mean ±standard error) compared to an SCC of 0.81 ±0.01 for the maximum entropy approach [Fig. 2D]. While the SCC for the GNN approach is lower than the maximum entropy approach, it is still higher than the SCC between experimental contact maps from different cell lines. The SCC between IMR90 and GM12878 contact maps of the same genomic regions is 0.58 ±0.02. This suggests that the GNN is capturing cell line specific patterns of genome structure. The SCC requires two user-defined parameters; we explore the effect of these parameters in Section S6 [Fig. S7].

The second metric is the HiC-Spector score [54]. HiC-Spector compares contact maps by treating them as graph Laplacian matrices. It computes the sum of Euclidean norms between the top *κ* Laplacian eigenvectors of the two Hi-C contact maps. The metric is rescaled to the range [0,1], where higher is better. The GNN approach has a HiC-Spector scores of 0.51 ± 0.02 compared to 0.49 ±0.02 for the maximum entropy approaches, which is not statistically significant by two-sided paired t-test.

We also include two simpler metrics, which both directly measure the similarity of genomic-distance normalized contact maps, 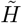. It is common to visualize the first principal component of 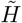, PC1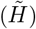, as a measure of A/B compartmentalization. For this reason, we compute the Pearson correlation between 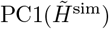 and 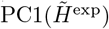 (Corr 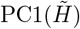 in Table 1). The GNN approach has a mean Pearson correlation of 0.90±0.01 between the first principal component of the simulated and experimental genomic-distance normalized contact maps. Meanwhile, the maximum entropy approach has a mean correlation of 0.78 ±0.03. Finally, we directly measure the root-mean-squared error between genomic-distance normalized contact maps. The GNN approach has an RMSE 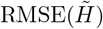 of 0.65 *±* 0.01 (vs 1.04 *±* 0.03 for maximum entropy).

We observe that the GNN approach performs better on the metrics computed from the genomic-distance normalized contact maps, 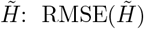 and Corr 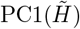 On the other hand, the maximum entropy approach performs better on the metrics computed from the raw contact maps, *H*: SCC and HiC-Spector. We discuss the possible origins and implications of this difference in Section S6.

### 2.3 Generalization to other Human and Mouse Cell Lines

For all results in this manuscript, we use a GNN trained on simulated data that was generated based on a single experimental Hi-C dataset of the IMR90 human cell line. To demonstrate that the GNN can generalize to other cell lines beyond IMR90, we use the GNN approach to simulate experimental contact maps corresponding to the even chromosomes of the human GM12878, HMEC, HUVEC, and HAP1 human cell lines. Further, to assess generalization to other organisms, we include the CH12.LX mouse cell line. Despite being trained exclusively on simulated data based on the IMR90 cell line, the simulated contact maps from the GNN approach reproduce experimental contact maps from these cell lines.

In Fig. 3A, we show example contact maps for the GM12878, HUVEC, and CH12.LX cell lines. The simulated contact maps reproduce A/B compartmentalization [Fig. 3B] and contact probability scaling [Fig. 3C]. In Fig. S2A, we show the simulated contact maps from the maximum entropy approach for comparison.

**Figure 3:**
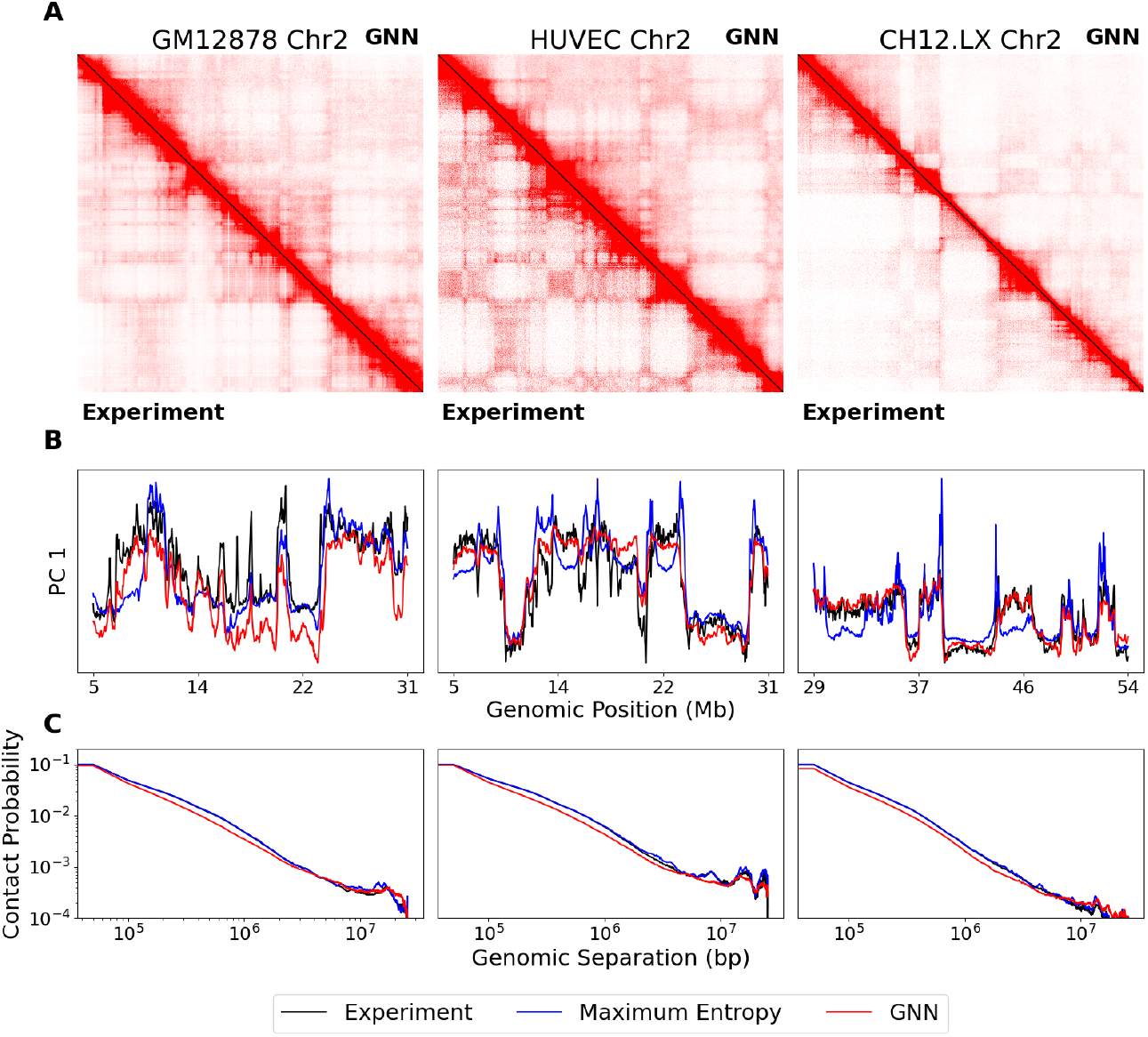
GNN generalizes to unseen cell types. *Left:* GM12878 human lymphoblastoid cell line Chr2:5.1-30.7 Mb. *Middle:* HUVEC murine umbilical vein endothelial cell line Chr2:5.1-30.7 Mb. *Right:* CH12.LX mouse lymphoma cell line. **A** Comparison of experimental and GNN simulated contact maps. The lower triangle shows the experiment, and the upper triangle shows the GNN simulation. The color bar is the same for all subfigures. SCCs between simulated and experimental contact maps are 0.67, 0.70, and 0.76, respectively. **B-C** Comparison of the experiment (black), maximum entropy (blue), and GNN (red). **B** Comparison of the first principal component of genomic-distance normalized contact maps. **C** Comparison of contact probability scaling.

Averaged over 158 simulated contact maps from the four human cell lines, the results are comparable to those seen on the IMR90 cell line [Table S3]. The GNN approach had an SCC of 0.71 ±0.01. In comparison, the maximum entropy approach had an SCC of 0.82 ±0.00. On all other metrics, the GNN approach outperforms the maximum entropy approach [Table S3]: HiC-Spector Score (0.54 ± 0.01 vs 0.52 ±0.01); Corr 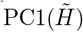 (0.85 ± 0.01 vs 0.79 ±0.01); and 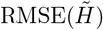 (0.70 ±0.01 vs 1.07 ±0.01). Further, the GNN approach was 6.6x faster on average.

The results for the mouse CH12.LX cell line are comparable to the results for the human cell lines. The GNN approach had an SCC of 0.78 ±0.01 compared to 0.81 ±0.01 for the maximum entropy approach, which is a smaller difference in SCC than for any of the human cell lines. The GNN approach outperforms the maximum entropy on all of the other metrics except for Corr 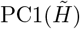 (0.80 ±0.04 vs 0.85 ±0.01) Table S3]. For the mouse CH12.LX cell line, the GNN approach was 8.3x faster on average.

Table S3 includes results for each cell line separately. We do not see substantial variation in performance between cell lines. These results demonstrate that the GNN has robustly learned properties of Hi-C data that generalize across cell lines.

### 2.4 Validation with Super-Resolution Imaging

To directly validate the structures estimated from simulations using GNN-estimated parameters, we compare our simulated structures with experimental imaging data. Given an experimental contact map of IMR90, we use the GNN to estimate the parameters of our polymer model, and we compare the estimated structures to super-resolution chromatin tracing experiments of IMR90 Chr2 [49] and Chr21 [4, 49].

We note that the energy functional of our simulation does not directly constrain distances, nor is the GNN trained to reproduce distances. The two parameters in our model that most directly affect distances are the bond length, *b*, between neighboring particles and the simulation volume, *V*. We choose a bond length (*b* = 200 nm) and simulation volume (*V* = 8 *µ*m^3^) such that the mean distance scaling as a function of genomic separation is comparable between the simulation and experiment [Section S2.1.5, Fig. 4AB, Fig. S4].

**Figure 4:**
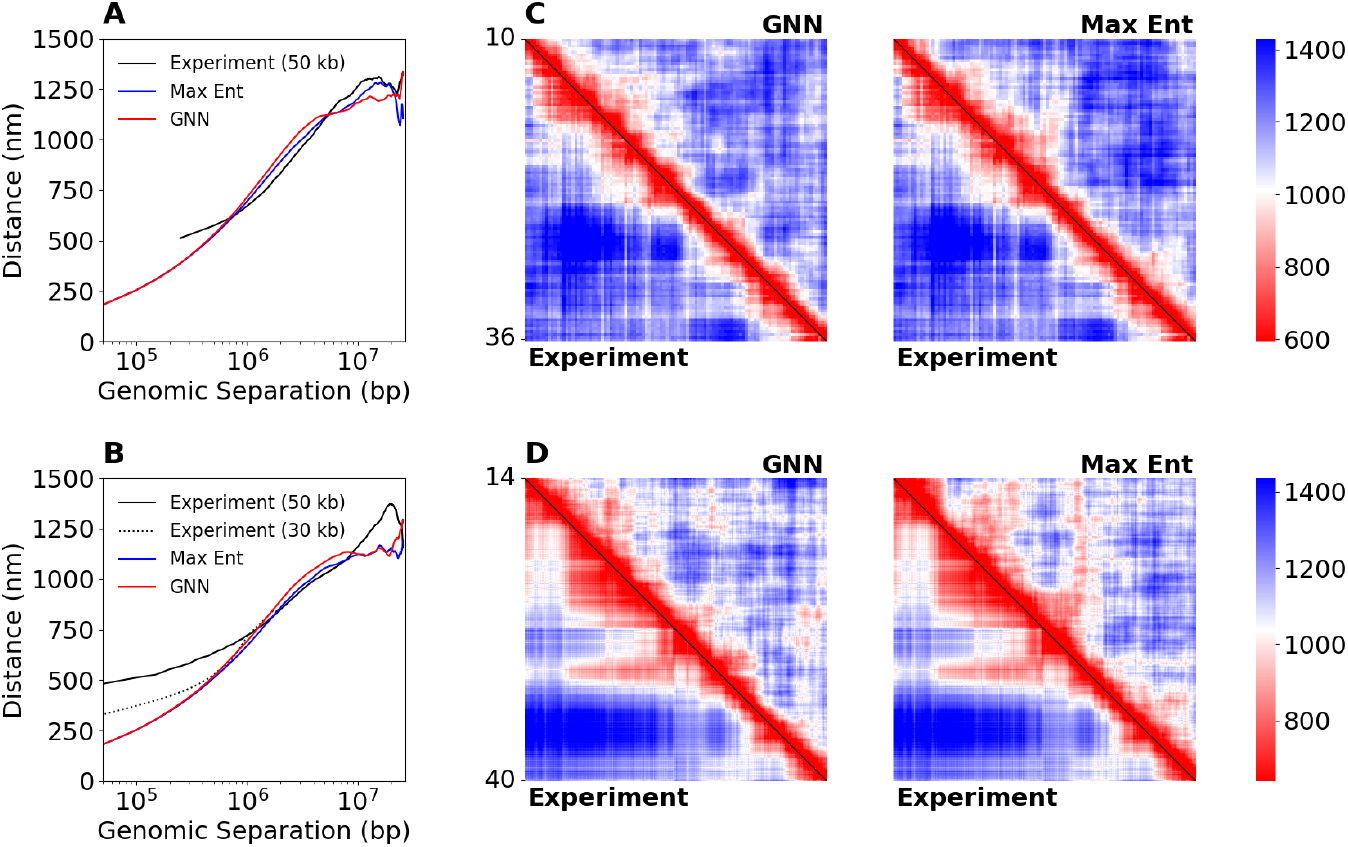
GNN reproduces patterns in spatial distances from super-resolution chromatin tracing experiments. We obtain experimental structures at 50kb resolution Su et al. [4, 49]. *Top row* : IMR90 Chr2:10-25.6 Mb. *Bottom row* : IMR90 Chr21:14-39.6 Mb. **A-B** Mean spatial distance scaling as a function of genomic separation for experiment (black), GNN simulation (red), and maximum entropy simulation (blue). **B** The additional dotted black line corresponds to an experimental structure of IMR90 Chr21:28Mb-29.2 Mb at 30 kb resolution from Bintu et al. [4]. **C-D** Mean spatial distance matrices. The lower triangle shows the experimental distance matrix. *Left:* The upper triangle shows the GNN simulated distance matrix. *Right:* The upper triangle shows the maximum entropy simulated distance matrix.

The simulated structures are in good agreement with the experimental structures. On chromosome 2, the simulated and experimental mean distance matrices have a Pearson correlation of 0.89 for the GNN (maximum entropy: 0.90) [Fig. 4C]. On chromosome 21, the simulated and experimental mean distance matrices have a Pearson correlation of 0.78 for the GNN (maximum entropy: 0.83) [Fig. 4D]. These results help suggest that the simulated structures are representative of the underlying experimental structures. The corresponding simulated contact maps are included in Fig. S3.

The accuracy of the simulated distances depends greatly on the choice of bonded parameters and bonded potential term. A more accurate bonded term in the energy functional could improve these results [21]. Importantly, the distance matrices from the GNN approach are comparable to the maximum entropy approach, as both methods use the same bonded term. Improving the underlying polymer model would improve both results.

## 3 Conclusion

To improve our understanding of genome organization and gene expression, characterizing chromatin structure is essential. With efficient approaches for modeling chromatin structure from Hi-C data, we can compare chromatin structures across many cells and cell types. These chromatin models can unveil the regulatory mechanisms behind gene expression to answer questions about cell differentiation and development.

We introduce an efficient computational pipeline for estimating chromatin structures from Hi-C data. Our approach combines advances in polymer modeling of chromatin structure with modern machine learning methods to predict chromatin structures significantly faster than classical approaches. A key contribution of our approach is that we train the GNN exclusively on simulated data. This was made possible by leveraging the polymer model to generate large quantities of simulated contact maps. Further, because we train the GNN to predict thermodynamic parameters of the polymer model instead of predicting structures, the GNN does not need to learn the physics of chromatin folding. We believe that this choice makes the GNN more robust to variation in genome structure across different cell lines as well as simplifies training. One obvious drawback of this choice is that we cannot overcome any limitations of the physics in our polymer model. As biophysical modeling of chromatin improves, we can retrain the GNN using more sophisticated chromatin models.

Using our GNN approach is an order of magnitude faster than using the maximum entropy approach to estimate the parameters of our polymer model. At the same time, the accuracy of the predicted chromatin structures is comparable to structures from simulations using parameters estimated with the maximum entropy approach. In addition, the GNN does not require *a priori* knowledge or assumptions about the particle labels, Ψ, of the chromatin model, whereas the maximum entropy approach requires the particle labels as an input. Notably, we show that despite training the GNN exclusively on cells from one human cell line (IMR90) it is still able to generalize to other human and even mouse cell lines.

Our approach has the potential to be useful in the analysis of single-cell variants of Hi-C [33]. These single-cell Hi-C variants can generate tens of thousands of single-cell contact maps, providing an opportunity to compare chromatin structures across cells and cell populations [14]. If using the existing maximum entropy approach, it would be incredibly computationally expensive to simulate the structure of each single-cell contact map. One challenge is the sparsity of single-cell Hi-C data poses a problem for both the GNN and maximum entropy approaches, as discussed in Section S3 [Fig. S5]. Future work will address how to generalize our graph neural network to the single-cell setting, potentially by leveraging existing imputation procedures such as scHiCluster [60] or Higashi [58] to infer missing contacts in single-cell Hi-C data *in silico*.

## 4 Materials and Methods

### 4.1 Experimental Data

We obtain experimental Hi-C contact maps, *H*, at 50 kb resolution. Experimental data files are downloaded from Juicebox [11] or ENCODE [17, 22, 29, 52], as indicated in Table S4. Contact maps are initially downloaded as raw counts without normalization. We include only autosomal chromosomes. We divide chromosomal contact maps into 25.6 Mb regions, corresponding to *m* = 512 bins per contact map. We only choose regions such that *H*_*ii*_ *>* 0 for all *i*. Doing so avoids centromeric and telomeric regions as well as regions with low read-mappability. We normalize the contact maps such that the average contact probability between beads *i* and beads *i ±* 1 is 0.1. We then overwrite the main diagonal (self-self interactions) with 1’s.

### 4.2 Particle-based Chromatin Model

We model chromatin as a heteropolymer of *m* particles, where each particle represents 50 kb of DNA. The potential energy function of our polymer model can be decomposed into bonded, 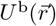, and nonbonded, 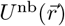, terms:

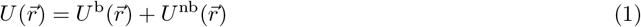

where 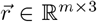 contains the positions of all *m* particles in the simulation and 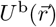 is a Gaussian chain bonded potential [Section S2.1].

The nonbonded term is derived from the Theoretically Informed Coarse Grained (TICG) potential for block copolymers [7, 8, 40]. While there are several existing chromatin models in the literature [9, 41, 47, 57], we choose the TICG potential because of the computational efficiency resulting from its grid-based implementation [Fig. 1B]. Instead of computing the spatial distance between particles, the TICG potential defines particles located in the same grid cell to be interacting 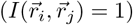. Omitting some constants, the nonbonded potential can be written as follows:

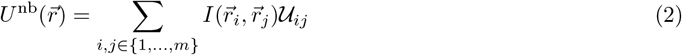

where 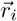 denotes the position of particle 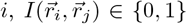 indicates if particles *i* and *j* are in same grid cell, and 𝒰 *∈* ℝ^*m×m*^ is a matrix of interaction parameters between pairs of particles.

The nonbonded potential can be further decomposed as:

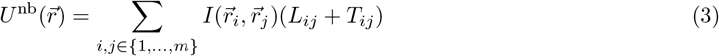

where *L, T ∈* ℝ^*m×m*^ are two types of interaction parameters and 𝒰 := *L* + *T*.

Note that the original description of the TICG potential is written in terms of local densities of particles within each grid cell rather than pairs of particles [8]. In Section S2.1, we include a description in the local density notation and a derivation of Eq. (2) and Eq. (3). We will use the notation in Eq. (2) and Eq. (3) for both the maximum entropy and GNN approaches.

Here, we will provide some intuition for the role of *L* and *T*. Both terms are analogous to terms in the MiChrom model proposed by di Pierro *et al*. [9]. *L* is low-rank by construction and controls interactions between particles of different epigenetic states. The *L* contribution is a generalization of the “chromatin types” term used in the MiChrom model. *T* is a symmetric matrix where all descending diagonals are constant-valued (i.e., a Toeplitz matrix). As a result, all particles with the same genomic distance, *d* = |*i−j*|, will have the same value *T*_*ij*_. *T* serves to modify the scaling behavior of the polymer chain to match the experimental contact probability scaling. The *T* contribution is analogous to the “ideal chromosome” term used in the MiChrom model [9]. We define 𝒰_*ij*_ := *L*_*ij*_ + *T*_*ij*_ as a net pairwise interaction strength. Intuitively, 𝒰 can be considered a look-up table for the interaction energy between every pair of particles *i* and *j*. Pairs of particles with negative 𝒰_*ij*_ will attract, while those with positive 𝒰_*ij*_ will repel.

Given 𝒰, we simulate chromatin structures by Monte Carlo sampling [Section S2.1.6]. With these structures, we can calculate any desired structural property. One property of particular interest is the simulated contact map, *H*^sim^, which we use to assess the quality of our simulation. We define *H*^sim^ as the ensemble average (i.e., averaged over all structures) of the grid indicator function, 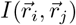:

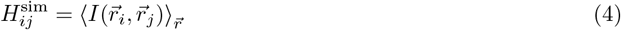

We typically estimate *H*^sim^ by averaging over 30,000 structures sampled via the polymer model. We expect the simulated contact map, *H*^sim^, to accurately reproduce the experimental contact map, *H*.

### 4.3 Data Generation

We obtain a set of experimental contact maps, *H*^(𝓁)^ for 𝓁 = 1, …, 33 corresponding to 25.6 Mb regions at 50 kb resolution of the odd chromosomes of the IMR90 cell line. Each contact map contains *m* = 512 bins (i.e., rows/cols). For each *H*^(𝓁)^, we use the maximum entropy approach to estimate 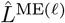 and 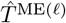 matrices from Eq. (3). Define 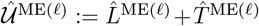. See Section S2.2 for a complete description of the maximum entropy optimization.

We use the following procedure to generate 10,000 synthetic interaction parameter matrices, 𝒰^(s)^ ∈*R*^*m×m*^ for s = 1, …, 10000:

I. Generate random *L*^(s)^ matrix [Algorithm S7]. *L*^(s)^ is constructed with random eigenvalues drawn from distributions estimated using kernel density estimation over the eigenvalues of 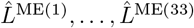. All 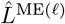 are rank 10, and therefore *L*^(s)^ is as well. The eigenspace of *L*^(s)^ is constructed from the eigenvectors of a random 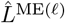.
II. Generate random Toeplitz matrix, *T*^(s)^. *T*^(s)^ is defined as equal to a random 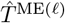 without any modification.
III. Define 𝒰 (s) = *L*^(s)^ + *T*^(s)^.

See Section S2.5 for further discussion.

We use our polymer model to simulate the corresponding contact map, *H*^(s)^, for each 𝒰^(s)^. As described in Section S2.1.6, we simulate chromatin structures via Metropolis Monte Carlo sampling. Each simulation results in 30,000 structures, and *H*^(s)^ is calculated from these structures according to Eq. (4). At the conclusion of the data generation procedure, we have 5000 pairs (*H*^(s)^, 𝒰^(s)^).

### 4.4 Neural Network

Generically, graph neural networks (GNNs) require three inputs, (*A, X, ε*). *A* is the adjacency matrix of the graph, *X* is node features, and *ϵ* is the edge features. We use the experimental Hi-C contact map to define all three of these aspects of the graph structure. We include an ablation study demonstrating the importance of the neural network design choices described here in Section S2.4.3 [Table S2].

We first obtain an experimental contact map, *H*^100kb^, binned at 100 kb resolution. While we simulate chromatin at 50 kb resolution, we use 100 kb resolution for the neural network as the GNN both trains faster and performs better when using a coarser resolution [Table S2]. We normalize the contact map by dividing by the mean of its main diagonal. Then we overwrite the main diagonal with 1’s.

We define *A*_*ij*_ = 1 if 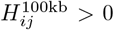 and *A*_*ij*_ = 0 otherwise. Since we use contact maps at 100 kb resolution, most values 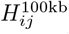 are non-zero, and the graph is nearly complete. If all values are non-zero, the graph will be complete (i.e., *A* will be a matrix of 1’s).

We define node features, *X*, as the top 10 eigenvectors of the 100 kb resolution genomic-distance normalized contact map, 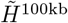.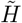 is calculated by dividing each entry of the contact map by the mean along the corresponding diagonal:

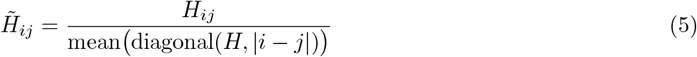

The eigenvectors of this matrix correlate well with biological features such as epigenetic modifications and chromatin compartmentalization [26, 43].

We define edge features between nodes *i* and *j* as:

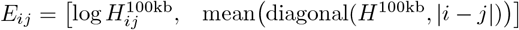

The first value, 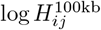, is the log of the corresponding entry of the 100 kb resolution contact map (*ϵ*_*ij*_ only needs to be computed when of *A*_*ij*_ = 1, so 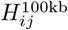 is guaranteed to be nonzero.). The second value, mean diagonal(*H*^100kb^, |*i − j*|), is the mean contact frequency between genomic loci at distance |*i − j*|, calculated by taking the mean along the |*i − j*|th diagonal of the experimental contact map.

Given these inputs, the GNN outputs 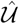. We train with mean squared error loss after a log-transformation of 𝒰 and 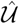, denoted as 𝒰^*†*^ and 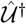. Specifically, we define 𝒰^*†*^ := sign(𝒰) ⊙ ln(|𝒰| + 1) where sign(𝒰) ∈ {−1, 1}^*m×m*^ contains the element-wise sign of 𝒰 and ⊙ is the Hadamard (element-wise) product. Computing the loss using the 𝒰^*†*^ representation improves empirical performance [Table S2].

Details of the training procedure are included in Section S2.4.2.

#### 4.4.1 Neural Network Architecture

Fig. S1 shows a schematic of the architecture. First, the node features are passed through a linear layer to form an initial node embedding (omitted in figure). The node embedding is updated by a series of graph message passing layers. The message passing architecture is a modification of the graph attention network from Brody *et al*. [5] [Section S2.4.1]. Between each message passing layer is a multi-layer perception (MLP) with leaky ReLU activations.

The resulting latent node embedding, *Z*, is further processed by two independent modules: an epigenetic module and a genomic-distance module. The epigenetic module computes a bilinear function of the node embedding, 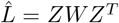, where *W* is a learnable weight matrix. 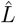 is then upsampled by a factor of two in order to map from 100 kb resolution to 50 kb resolution. The genomic-distance module is an MLP that acts on the flattened node embedding. The output of this MLP is used to construct 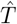 (at 50 kb resolution). We combine the two modules by adding their results (element-wise) to define 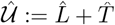.

Finally, to account for the sign-invariance of the eigenvector node features, we adopt an idea from the SignNet architecture proposed by Lim *et al*. [27]. The entire GNN architecture is run twice, once with node features defined by positive eigenvectors, *υ*, and once by the negated eigenvectors, −*υ*, to account for the sign ambiguity of the eigendecomposition. The two resulting predictions are summed to yield a final prediction. We omit this step from Fig. S1 for simplicity of presentation.

### 4.5 Code Availability

The source code for implementing, training, and applying our approach is available at the GitHub repository: https://github.com/ERSchultz/GNN_HiC_to_Structure.git.

## S1 Pseudocode

Algorithm S1 defines our simulation procedure. See Section S2.1.6 for a description of the Monte Carlo sampling procedure. We use this simulation procedure for both the maximum entropy approach [Algorithm S4] and the GNN approach [Algorithm S5]. Algorithm S2 defines how we generate simulated contact maps. Algorithm S4 defines the maxim entropy optimization approach for estimating simulation parameters. Line 7 calls our Newton’s method procedure, which we don’t define for simplicity. See Section S2.3.4 for details of the maximum entropy optimization. Algorithm S5 defines the GNN approach for estimating simulation parameters. Line 1 calls our graph neural network, which we don’t define for simplicity.

### Algorithm S1

*H*^sim^ ← SimulationEngine

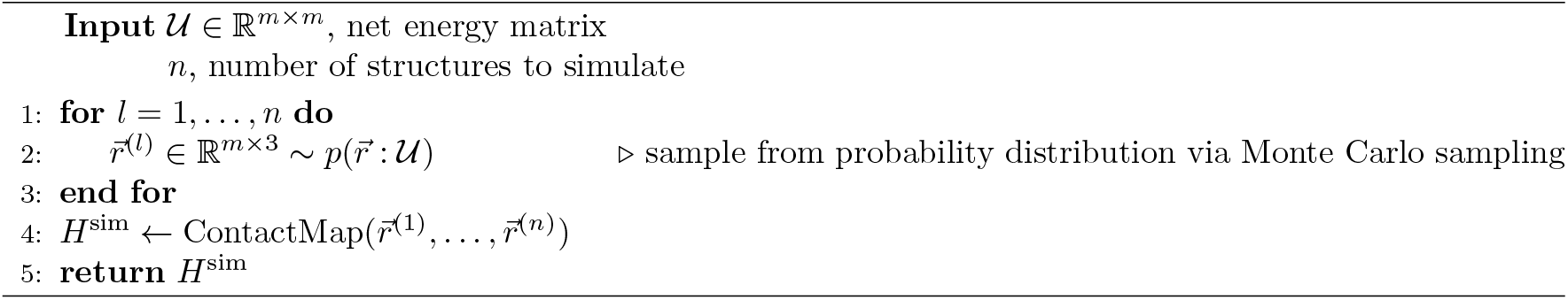

### Algorithm S2

*H*^sim^ ← ContactMap

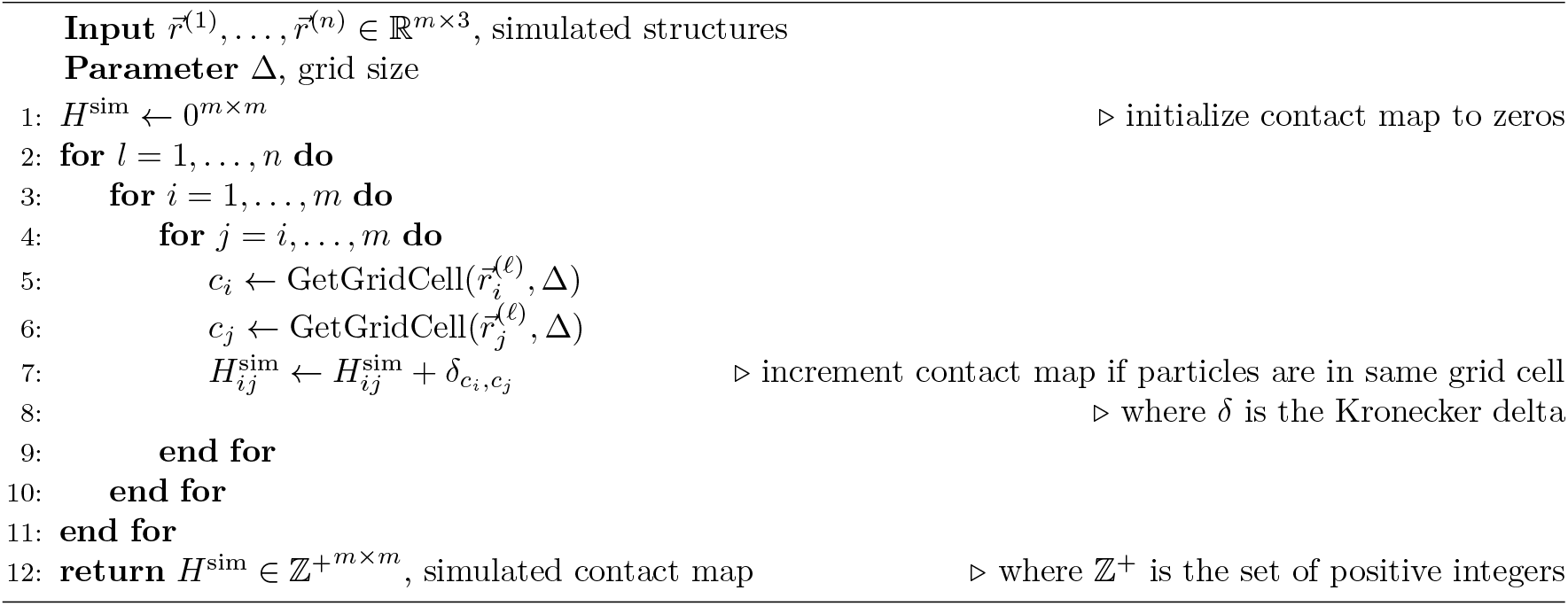

### Algorithm S3

*c* ← GetGridCell

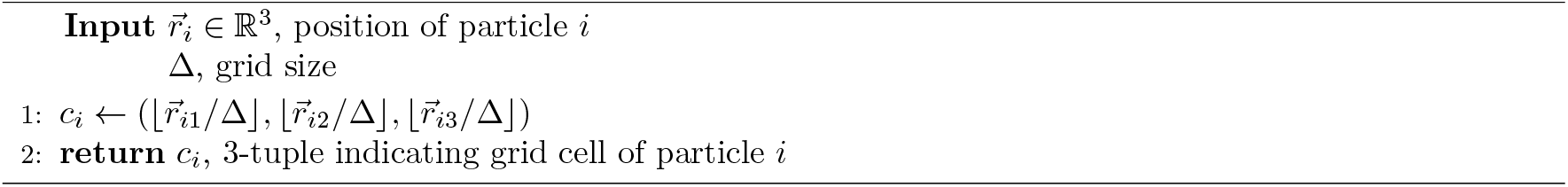

### Algorithm S4

*H*^sim^ ← MaximumEntropyOptimizationApproach

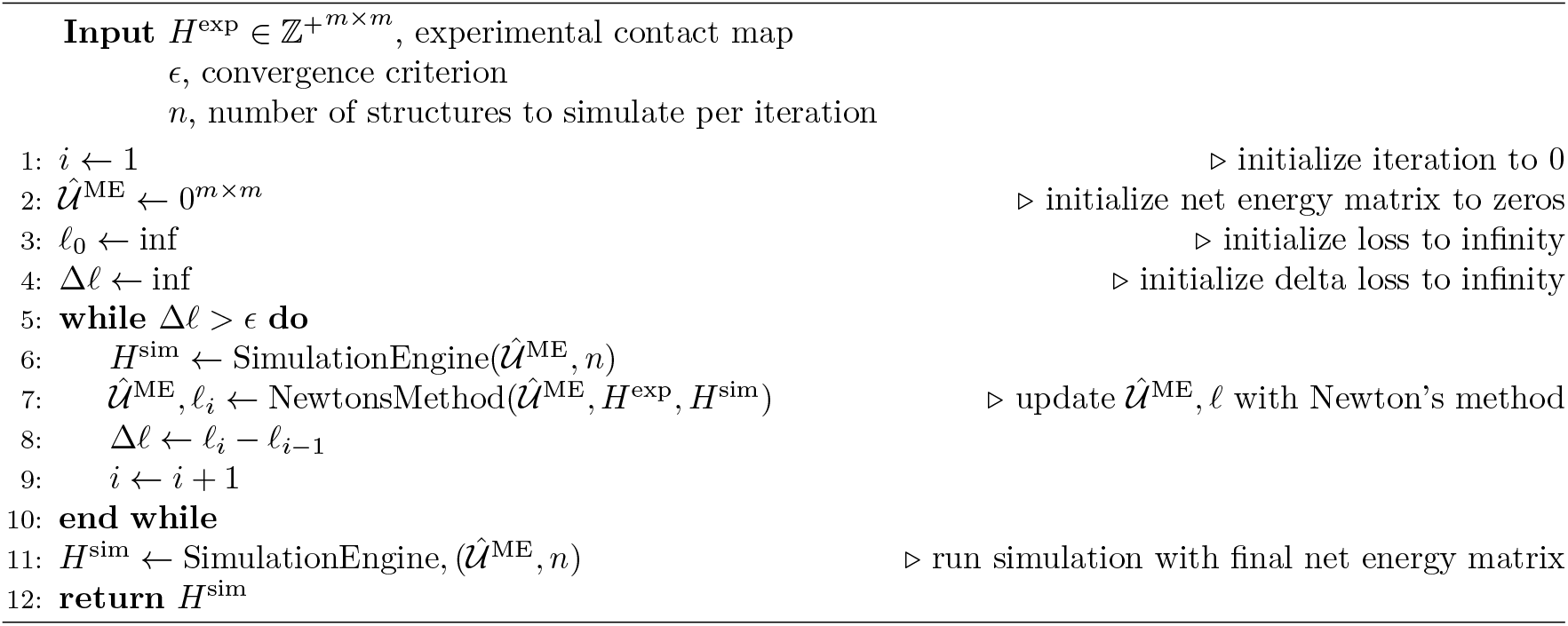

### Algorithm S5

*H*^sim^ ← GNNApproach

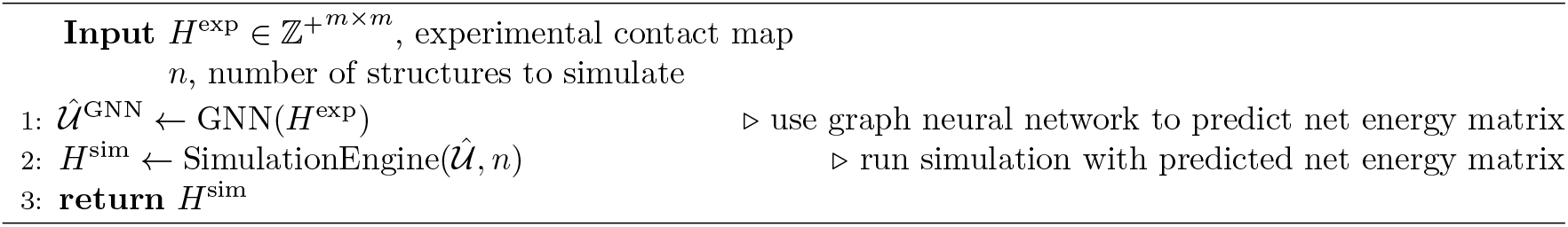

## S2 Supplementary Methods

### S2.1 Polymer Model Details

Similar to other top-down approaches for chromatin modeling [9, 41, 47, 57], we employ a generic coarse-grain heteropolymer model as the basis for our polymer model. Then, we build in additional complexity using experimental data as constraints in a top-down fashion. We treat 50 kb of DNA as one simulation particle. The energy functional of our polymer model can be decomposed as a sum of bonded, 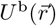, and nonbonded, 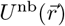, contributions:

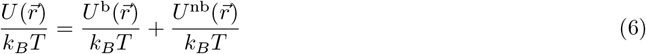

where 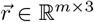 contains the positions of all *m* particles in the polymer, *k*_*B*_ is the Boltzmann constant, and *T* is temperature.

In a slight abuse of notation, we omit 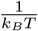 from the remainder of the equations in this manuscript for simplicity.

The bonded interaction controls the interaction between particles *i* and *i* + 1. We use a Gaussian chain bonded interaction defined as follows:

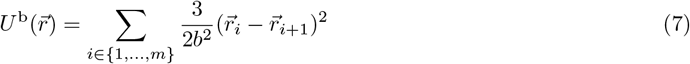

where *b* is the desired mean bond length and 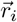 is the position of the *i*th particle.

The non-bonded interaction describes the interaction between pairs of particles and can be further decom-posed into an epigenetic, 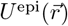, and genomic distance, 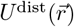, contribution.

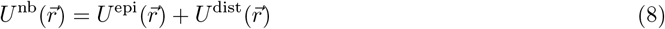

#### S2.1.1 Epigenetic Contribution

The epigenetic contribution reproduces phase separation between particles with different chromatin states. For now, consider each particle to be labeled with a mutually exclusive chromatin state label. In particular, we use the Theoretically Informed Coarse Grain (TICG) potential [7, 8, 40]. The TICG potential is a meanfield technique originally developed to study polymer phase separation. Notably, the TICG potential has previously been used to model chromatin structure, although in a bottom-up fashion [30].

The TICG potential calculates nonbonded interactions via a field-theoretic Hamiltonian based on the local density of the polymer calculated upon a grid-based discretization of cartesian space.

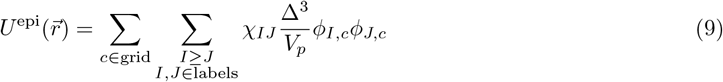

where *χ*_*IJ*_ is the Flory-Huggins interaction energy between particles with label *I* and with label *J, ϕ*_*I,c*_ is the volume fraction of particles with label *I* in grid cell *c* for a given configuration 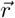, Δ is the grid size, *V*_*p*_ is the volume of an individual particle.

The advantage of the TICG functional is that the structural ensemble can be efficiently sampled via Monte Carlo sampling (detailed in Section S2.1.6) in 𝒪 (*m*) time where *m* is the number of particles. As a result, our simulation is sufficiently efficient to enable the simulation of entire chromosomes at nucleosome resolution (200 bp) [30]. However, in this work, we use a resolution of 50 kb as a proof of concept of our approach.

#### S2.1.2 Genomic Distance Contribution

The genomic distance contribution reproduces the contact probability scaling seen in experimental Hi-C data. This term is inspired by and analogous to the “ideal chromosome” term in the MiChrom introduced by Di Pierro *et al*. [9]. By design, this term is invariant to translation along the genome and is independent of the epigenetic contribution.

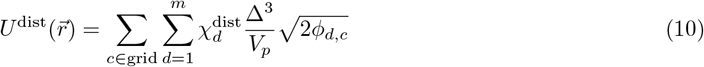

where *d* = |*i− j*| is the distance along the polymer chain between particles *i* and *j*, 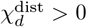 is the interaction energy between particles with genomic separation *d, ϕ*_*d,c*_ is the volume fraction of *bonds* between particles of genomic distance *d* in grid cell *c* for a given configuration 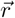.

#### S2.1.3 Net Interaction Representation

To make the machine learning task more tractable we re-write *U* ^nb^. A brief outline of the derivation is shown here, and a full derivation is included below in Section S2.1.4.

The epigenetic contribution, *U* ^epi^, can be rewritten as:

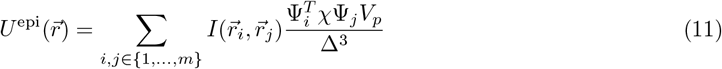

where 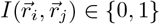 indicates if particle *i* and *j* are in the same grid cell, Ψ_*i*_ = {0, 1}^*k*^ is a binary vector containing the labels of particle *i*, and *χ* is an upper triangular matrix containing every *χ*_*I,J*_

In contrast to Eq. (9), which was a double sum over all grid cells and all labels, Eq. (11) is a sum over all pairs of beads. In Eq. (9), the particles have discrete labels, and therefore Ψ_*i*_ is a binary vector. However, Ψ_*i*_ can be real-valued in general. For convenience, we define 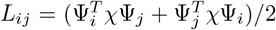. *L*_*ij*_ contains the net epigenetic interaction energy between particles *i* and *j* and is symmetric by construction. *L* can be factorized to recover *χ* if the particle labels, Ψ, are known.

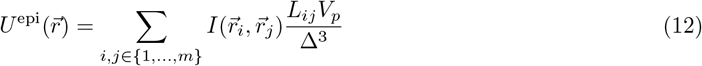

where *L*_*ij*_ is the net epigenetic interaction between particles *i* and *j*.

By a similar approach, the genomic distance contribution, *U* ^dist^, can be rewritten as:

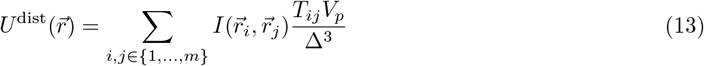

where 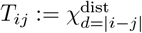 is the genomic distance interaction between particles *i* and *j*.

By construction, *T* has a main diagonal of zeros, and all descending diagonals are constant-valued. Such a matrix is referred to as a Toeplitz matrix.

We can combine the genomic distance (13) and epigenetic contributions (12) into a single non-bonded term:

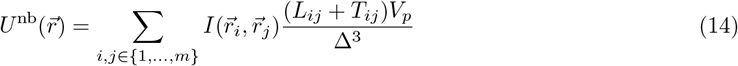

#### S2.1.4 Derivation

Consider Eq. (9):

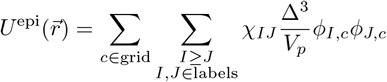

where *χ*_*IJ*_ is the Flory-Huggins interaction energy between particles with label *I* and with label *J, ϕ*_*I,c*_ is the volume fraction of particles with label *I* in grid cell *c* for a given configuration 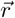, Δ is the grid size, and *V*_*p*_ is the volume of an individual particle.

By definition:

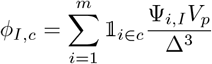

where 𝟙_*i∈c*_ = 1 if particle *i* is in grid cell *c* (0 otherwise) and Ψ_*i,I*_ = 1 if particle *i* has label *I* (0 otherwise). By substituting *ϕ*_*I,c*_:

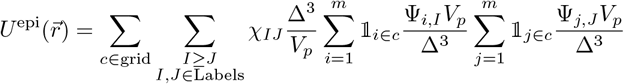

Which can be rewritten as:

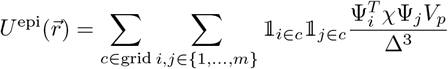

where ∑_*i,j∈*{1,…, *m*}_ is a sum over all pairs of particles (*i, j*) and *χ* is an upper triangular matrix containing every *χ*_*I,J*_.

Which can be rewritten as Eq. (11):

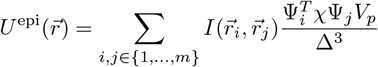

where 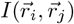 indicates if particle *i* and *j* are in the same grid cell.

Define 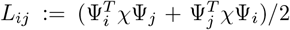. Then, *L*_*ij*_ contains the net epigenetic interaction energy between particles *i* and *j* and is symmetric by construction [Eq. (12)]:

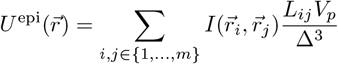

where *L*_*ij*_ is the net epigenetic interaction between particles *i* and *j*.

Now consider Eq. (10):

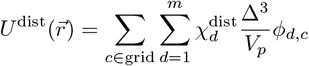

where *d* = |*i− j*| is the distance along the polymer chain between particles *i* and *j*, 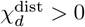 is the interaction energy between particles with genomic separation *d*, and *ϕ*_*d,c*_ is the volume fraction of *bonds* between particles of genomic distance *d* in grid cell *c*.

By definition:

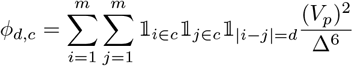

By substitution, the following is true:

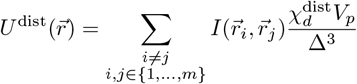

We can rewrite this equation by defining 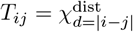 and *T*_*ii*_ = 0 [Eq. (13)]:

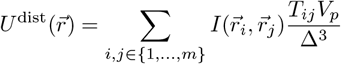

Finally, we can clearly see that the genomic-distance dependent [Eq. (13)] and epigenetic effects [Eq. (12)] can be combined as follows into a single non-bonded term [Eq. (14)]:

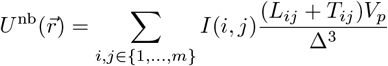

Alternatively,

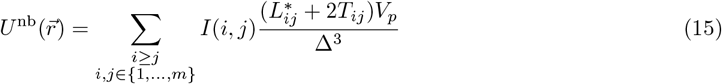

where *L*^*∗*^ = 2*L − I*_*m*_ ⊙ (*L*), where ⊙ is the Hadamard product and *I*_*m*_ is the size *m* identity matrix. Define 𝒰 = *L* + *T* and 𝒰^*∗*^ = *L*^*∗*^ + 2*T*. Eq. (15) is more efficient than Eq. (14) since it only considers pairs of particles where *i* ≥ *j*. For this reason, we use Eq. (15) during the Monte Carlo simulation.

#### S2.1.5 User-Defined Parameters

All simulations described in the main text use a particle volume, *V*_*p*_, of 130,000 nm^3^, a bond length, *b*, of 200 nm, a simulation volume, *V*, of 8 *µ*m^3^, and the simulation boundary is a prolate spheroid with an aspect ratio of 1.5. The GNN is trained on a simulated dataset using these bonded parameters, and therefore, it may not extrapolate to other choices of bonded parameters. However, the GNN does not depend in any way on the choice of bonded parameter; we could train a GNN on simulated data using any reasonable choice of bonded parameters.

The particle volume, *V*_*p*_, is defined based on the resolution of the simulated chromatin fiber. A coarse-grained particle at nucleosome resolution (200 bp per particle) is assumed to have a particle volume of 520 nm^3^ as in the polymer model of MacPherson *et al*. [30]. Since we use 50 kb resolution for all simulations, we rescale the particle volume using a Gaussian renormalization [44], resulting in a particle volume of 130,000 nm^3^.

We choose the bond length, *b*, simulation volume, *V*, and aspect ratio based on the experimental distance scaling [Fig. S4] from Su et al. [49]. Using (*b* = 200 nm, *V* = 8 *µ*m^3^, aspect ratio = 1.5) yields the best reproduction of the distance scaling as a function of genomic distance. Interestingly, we find that a prolate boundary better reproduces experimental distance scaling than a spherical boundary [Fig. S4C]. This result is consistent with experimental evidence that chromosome structures are aspherical [23].

The grid size, Δ, is chosen such that a simulation with only bonded interactions matches the average experimental contact frequency between adjacent particles (*i* and *i ±* 1). Since all experimental contact maps are normalized such that the average contact frequency between adjacent particles is 0.1, we use the same grid size for all simulations (Δ = 151.6 nm).

Within the maximum entropy approach, the particle labels, Ψ in Eq. (11), are user-defined parameters. Prior works choose the labels to represent ‘chromatin states’, which are binary and mutually exclusive [9, 41]. The default TICG epigenetic term [Eq. (9)] assumes that particles have binary labels but does not require that these labels be mutually exclusive. Further, rewriting the epigenetic term as in Eq. (11) enables us to use real-valued labels. We use the first *k* = 10 principal components of the genomic-distance normalized contact map, 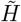, as the particle labels. We choose *k* = 10 because the simulation performance begins to plateau for *k >* 10 when using the maximum entropy approach [Fig. S6]. Note that the GNN does not require Ψ as an input, and therefore does not depend on *k*. However, for ease of comparison to the maximum entropy approach, we only train the GNN on simulated data where the epigenetic interaction matrix, *L*, is rank *k* = 10.

#### S2.1.6 Monte Carlo Simulation

We sample the equilibrium ensemble of our polymer model with Metropolis Monte Carlo sampling. Simulations are run for 300,000 Monte Carlo cycles. We employ four Monte Carlo moves: 1) Displacement, 2) Translation, 3) Crankshaft, and 4) Pivot. A Monte Carlo cycle consists of *m* Displacement, 500 Translation, 500 Crankshaft, and 50 Pivot moves, where *m* is the number of particles. We record the current configuration every 10 cycles, resulting in 30,000 structures per simulation.

Displacement translates a single random particle by 5 nm in a random direction.

Translation displaces a random polymer segment by 30% of the bond length in a random direction. The random polymer segment is defined by first choosing a random particle, then by choosing a second random particle from a two-sided exponential distribution with scale parameter 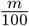.

Crankshaft rotates a random polymer segment about the axis defined by its ends. The random polymer segment is chosen as in Translation, except the polymer segment cannot contain the first or last bead. The polymer segment is then rotated by a random rotation in the interval 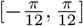radians.

Pivot rotates one end of the polymer about a random axis. One of the two polymer ends is chosen randomly, and a random particle is chosen. The resulting polymer end segment is rotated by a random rotation in the interval 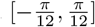radians around a randomly chosen axis.

In addition to these moves, we reposition the simulation grid during each Monte Carlo cycle. This prevents the discrete grid implementation from causing simulation artifacts at grid boundaries.

If any Monte Carlo move causes any specific grid cell to have a chromatin volume fraction greater than 0.5, we reject that move.

### S2.2 Maximum Entropy Optimization

### S2.3 Overview

The values of *χ*_*IJ*_ in Eq. (9) and 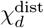 in Eq. (10) are not known *a priori* and must be inferred from experimental data. The maximum entropy principle is a common optimization framework for inferring unknown parameters from experimental data [18]. This approach has been applied previously to chromatin simulations (reviewed in [28]) and we use it as a baseline for comparison to our neural network-based approach.

In the maximum entropy approach, a model, *U* ^ME^, is learned as a correction to a more naive model. In our polymer model, we define 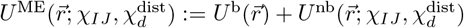. The homopolymer bonded term, *U* ^b^, is the naive model, and the nonbonded term, *U* ^nb^, is the maximum entropy correction term.

We seek to estimate *χ*_*IJ*_ and 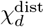 such that user-defined experimental observables, 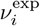, match their simulated counterparts:

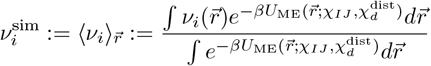

We define our experimental observables based on the experimental contact map (Section S2.3.2). We will optimize *χ*_*IJ*_ and 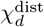subject to the constraint that 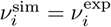 (Section S2.3.4). Note that the constraints are not guaranteed to hold with equality. Specifically, we seek to maximize the information entropy subject to our constraints:

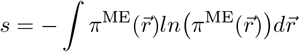

where *π*^ME^ is the maximum entropy probability density.

Our description of the maximum entropy approach is adapted from the supplement of di Pierro *et al*. [9].

#### S2.3.1 Notation

*m* is the number of particles in the simulation

*H*^exp^ *∈* ℤ^+*m×m*^ is an experimental Hi-C contact map (ℤ^+^ is the set of positive integers)

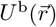 is a simple homopolymer (bonded) energy functional - we use a Gaussian chain potential

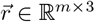 contains the coordinates of all m particles in 3D space

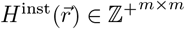 is a *instance* simulated contact map (i.e., corresponding to a single configuration *r*)

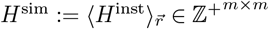 is the *ensemble* simulated contact map (i.e., the ensemble average over *r*)

Ψ ∈ ℝ^*m×k*^ are the particle labels

*ψ*_:, *I*_ ∈ ℝ^*m*^ is a column in Ψ

#### S2.3.2 ME Problem Formulation

Given *H*^exp^, 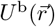, and Ψ the maximum entropy procedure seeks a modification of 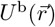 such that *H*^sim^ approximates *H*^exp^.

More explicitly, the maximum entropy (ME) procedure seeks to find a probability distribution (*π*^ME^) such that the following constraints hold:

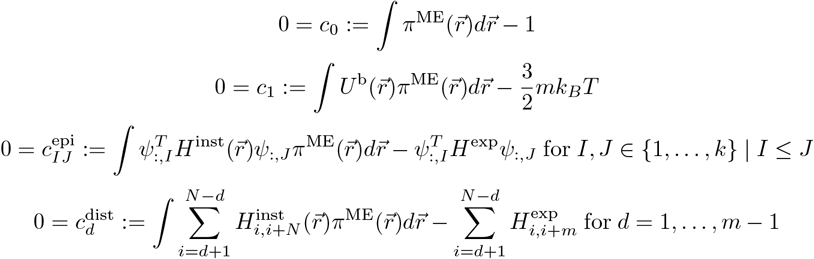

where *c*_0_ constrains the distribution to be normalized, *c*_1_ constrains the average potential energy to be equal to the thermal energy (where *k*_*B*_ is the Boltzmann constant and T is temperature), 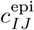 constrains the average contact frequency between beads of type *I* and type *J* to be the same in the experiment and the simulation, and 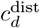 constrains the average contact frequency along each diagonal to be the same in the experiment and the simulation (in practice, we typically coarse-grain 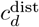 from *d* constraints to 96 constraints).

#### S2.3.3 Derivation of ME Probability Distribution

To optimize the parameters of *π*^ME^, we seek to maximize the information entropy subject to our constraints:

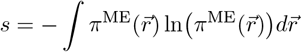

Using Lagrange multipliers we obtain the following extremum condition:

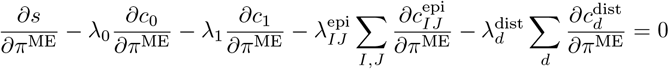

where each Lagrange multiplier, *λ*, corresponds to one of the constraints defined above.

Omitting the intermediary steps of the derivation, and renaming the following Lagrange multipliers:

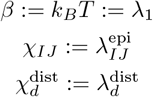

We arrive at the following probability distribution:

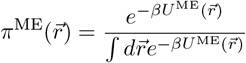

In our work, the resulting maximum entropy energy functional is:

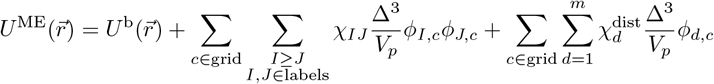

#### S2.3.4 ME Optimization Procedure

For the generic observable 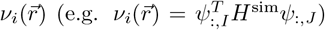 we wish to determine the parameters *λ*_*i*_ (e.g. *χ*_*IJ*_) contained in the potential energy:

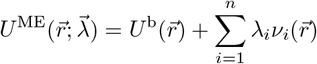

such that the expectation values of each observable coincides with its experimental value:

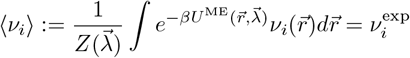

where 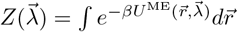 is the partition function.

We can define a convex objective function *θ*:

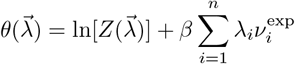

The partial derivatives of the target function are:

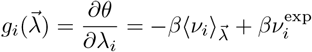

The Hessian is:

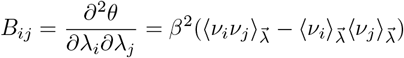

To find the optimal set of parameters, 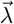, we numerically minimize the target function *θ* using Newton’s method:

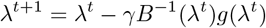

where *t* is the iteration number and *γ* is a dampening parameter.

We use *γ* = 0.25 for the first two iterations and *γ* = 1 for all subsequent iterations.

We define convergence when the normalized magnitude of the gradient is less than a convergence criterion, *ϵ*:

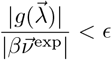

where 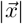 denotes the *l*2-norm of 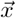

### S2.4 Neural Network Details

#### S2.4.1 Message Passing Architecture

Our message passing architecture is adapted from the graph attention network proposed by Brody *et al*. [5]. Compared to Brody *et al*., we include the edge features during the message passing step in addition to when calculating the attention coefficients.

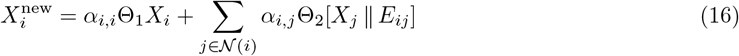

where 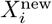 is the updated node embedding for node *i*, Θ_1_ and Θ_2_ are learnable weight matrices, 𝒩 (*i*) is the set of neighbors of node *i*, denotes concatenation, *E*_*ij*_ is the edge features for the edge between node *i* and *j*, and *α*_*i,j*_ is an attention coefficient defined as in Brody *et al*.:

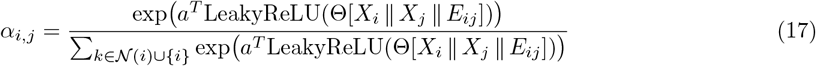

where *a* and Θ are learnable weight matrices.

#### S2.4.2 Neural Network Training

All neural networks are implemented and trained in PyTorch [38] using the PyTorch Geometric library [13]. We train/test on a synthetic dataset of *N* = 10000 samples generated as described in Section 4.3. We train on 9000 samples and test on the remaining 1000.

We use the Adam optimizer with an initial learning rate (lr) of 10^−4^ and other parameters set to their defaults. We train for 40 epochs at lr=10^−4^ followed by another 20 epochs with lr=10^−5^.

#### S2.4.3 Ablation Study

To assess the impact of variants of our data generation and training procedure, we performed an ablation study [Table S2]. We include all four metrics discussed in the main text in Table S2: SCC, HiC-Spector, Corr PC1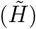, and RMSE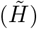 For simplicity, we only discuss the SCC here.

*Baseline* refers to the GNN as described in the main text. All results in the main text use the Baseline GNN.

*No log-transform in loss* is a GNN trained using the mean-squared error between 𝒰 and 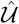 instead of between 𝒰^*†*^ and 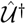. Training the GNN in this way noticeably decreases the SCC (0.573 vs 0.725), motivating our choice of defining the loss in the log space.

*50 kb resolution* uses a contact map at 50 kb instead of 100 kb resolution to define the graph structure. While this variant achieves lower MSE loss on the simulated validation set (0.313 vs 0.406), the SCC on the experimental validation set is slightly lower (0.650 vs 0.725). This result suggests that the GNN may be overfitting to the simulated contact maps when using the finer 50kb resolutione.

*Without SignNet* omits the SignNet step described in Section 4.4.1, where the entire GNN architecture is run twice, once with node features defined by positive eigenvectors and once by negative eigenvectors. We consider two variations. One where the eigenvectors are still included as a node feature, but their sign is unaccounted for - *Without SignNet (with eigenvectors)*. This variant performs substantially worse than the baseline (SCC 0.462 vs 0.725). And a second variation where the eigenvectors are omitted as well - *Without SignNet (without eigenvectors)*. Instead, the node features are initialized as a single constant value. Interestingly, this variant performs better than *Without SignNet (with eigenvectors)*, where the eigenvectors are included naively (SCC 0.647 vs 0.462).

*Without* mean(diagonal(*H*, |*i ™ j* |)) *in ε*_*ij*_ omits mean(diagonal(*H*, |*i ™ j* |)) from the edge features. This variant performs substantially worse (SCC 0.457 vs 0.725). Intuitively, we believe this feature to be important for the GNN to incorporate information about the contact probability scaling of the input contact map.

*Original message passing layer from [5]* uses the message passing layer as defined in Brody et al., instead of the modification we propose in Eq. (16). This variant performs dramatically worse (SCC −0.003 vs 0.725), demonstrating that incorporating edge features into the message passing step directly (not just when computation attention coefficients) is an essential modification.

### S2.5 Data Generation Details

#### S2.5.1 Motivation

A major challenge in our problem setting is obtaining sufficient training data. To train the GNN in a supervised manner, we require a large dataset of contact maps, *H*, and the corresponding parameters, 𝒰. There are two challenges to acquiring such a dataset. First, the number of existing experimental Hi-C contact maps is quite small compared to the scale of data needed to train modern neural networks. For instance, the ENCODE database contains only 195 different Hi-C experiments [29, 52]. Second, we do not know the corresponding polymer model parameters for these experimental contact maps, and estimating them via the maximum entropy approach would be computationally demanding. To address these challenges, we use synthetic data instead. The generalization properties and performance of the resulting trained neural network depend heavily on how the synthetic data is generated.

We seek to generate a training dataset of (𝒰, *H*) pairs. Given a synthetic parameter matrix, 𝒰^s^, we can use our polymer model to produce the corresponding simulated contact map, *H*^sim^. The challenge is to generate many 𝒰^s^ that are both biologically realistic and sufficiently diverse to allow the trained network to generalize well. We wish to leverage existing experimental data in combination with maximum entropy optimization solutions to strategically generate synthetic parameters. By using the maximum entropy parameter estimates to guide how we choose 𝒰^syn^, we can create realistic synthetic parameters. We expect these realistic parameters to yield simulated contact maps that resemble the experimental contact maps.

#### S2.5.2 Method

First, we obtain a set of experimental contact maps at 50 kb resolution, *H*^(𝓁)^ for 𝓁 = 1, …, 33 corresponding to 25.6 Mb regions of the odd chromosomes of the IMR90 cell line. Each contact map contains *m* = 512 bins (i.e., rows/cols). For each *H*^(𝓁)^, we use the polymer model with the maximum entropy approach to estimate 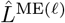 and 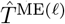 matrices from Eq. (3). Define 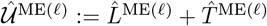. We will use these maximum entropy energy matrices to guide the generation of synthetic data.

We generate 10,000 synthetic interaction parameter matrices, 𝒰^(s)^ ∈ *R*^*m×m*^ for s = 1, …, 10000 using the procedure in Algorithm S6. In Algorithm S6, we first generate *L*^(s)^ using Algorithm S7. Next, we generate *T*^(s)^ as equal to the T matrix from a random maximum entropy simulation, 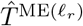. We find that this simple procedure for defining *T*^(s)^ is sufficient; the generative procedure for *L*^(s)^ in Algorithm S7 helps generate diverse simulated contact maps.

To facilitate generating random *L*^(s)^ matrices with Algorithm S7, we first fit a distribution to the eigen-values of 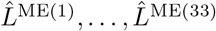. For each 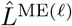, we compute the rank 10 eigendecomposition as 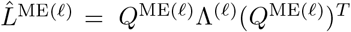 *Q*^ME(𝓁)^Λ^(𝓁)^(*Q*^ME(𝓁)^)^*T*^ where *Q*^ME(𝓁)^ *∈* ℝ^*m×*10^ contains the eigenvectors of 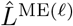 and 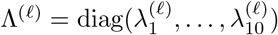 is a diagonal matrix containing the eigenvalues of 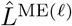. Note that 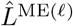 is rank 10 by construction [Section S2.1.5], so this eigendecomposition is exact. Next, we fit a probability distribution, *p*(*λ*_*j*_) for *j* = 1, …, 10, by kernel density estimation (KDE) over 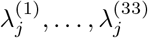. For the KDE, we use a Gaussian kernel with bandwidth 10 using the implementation in Scikit-learn [39]. When fitting the KDE, we ignore outliers as defined by 1.5 times the interquartile range.

In Algorithm S7, we define *L*^(s)^ based on its eigendecomposition. First, we generate random eigenvalues, 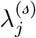, by drawing from the probability distributions *p*(*λ*_*j*_) (lines 1-2). Next, we define the eigenvectors, *Q*^(s)^, as equal to the eigenvectors from a random maximum entropy simulation, 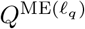 (line 3). In principle, we could design a generative model to generate random eigenvectors. However, we find using the maximum entropy eigenvectors without modification to yield sufficient results.

As an alternative to the approach in Algorithm S7, we could generate *L*^(s)^ by generating synthetic bead labels, Ψ ∈ ℝ^*m×k*^, and Flory-Huggin’s parameters, Ψ ∈ ℝ^*k×k*^ [Section S2.1][Eq. (11)]. Note that *L* := (Ψ*χ*Ψ^*T*^ + (Ψ*χ*Ψ^*T*^)^*T*^)*/*2. In our approach, we only need to generate *k* eigenvalues. To generate a synthetic *χ* would require 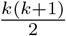 values (*χ* is symmetric). Further, the ordering of the bead labels in Ψ is ambiguous. Consider that any column permutation of Ψ is equally valid given the appropriate permutation of *χ*. In our approach, the eigendecomposition naturally orders the eigenvectors by their eigenvalue, which makes generating synthetic eigenvalues more straightforward.

##### Algorithm S6

𝒰^(s)^ ← Generate𝒰

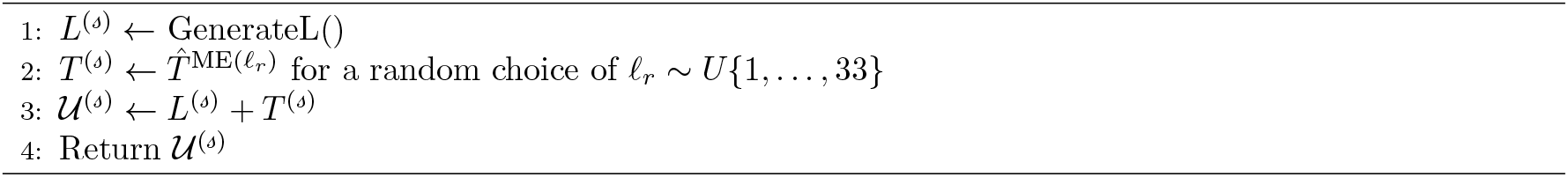

##### Algorithm S7

*L*^(s)^ ← GenerateL

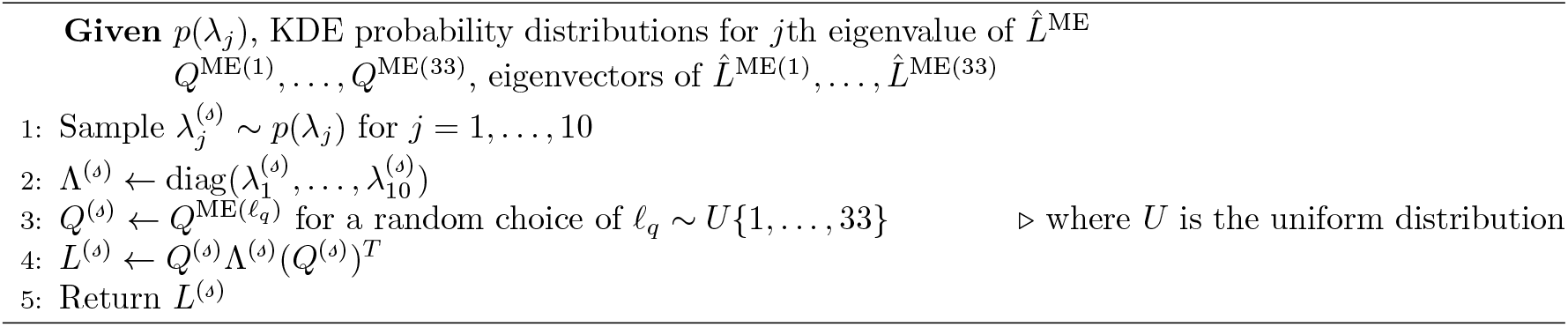

## S3 Robustness to Hi-C data quality

To demonstrate that the GNN is robust to the quality of the experimental data, we simulate the same genomic region using target contact maps with different numbers of total contacts. Contact maps with greater total contacts are of higher quality simply due to the law of large numbers. In our simulation, we can increase the number of total contacts by running for a random choice oflonger simulations and sampling more structures. In the experiment, the total number of contacts is equal to the number of sequencing read-pairs (i.e., sequencing read depth) used to generate the contact map. Deeper sequencing runs will increase the number of total contacts.

To systematically achieve an *in silico* contact map with a desired number of total contacts, we randomly sample contacts with probability proportional to the contact probability from a high-quality experimental contact map until we reach the desired number of total contacts. We define the total contacts as the number of contacts in the target genomic region (as compared to genome-wide). Fig. S5A-B shows the resulting *in silico* contact maps and corresponding simulated contact maps for a 25.6 Mb region of IMR90 Chr2 at 10^5^, 10^6^, and 10^7^ total contacts. The original experiment had ∼1.7 *×* 10^7^ contacts in this region.

We performed this sampling procedure for ten different IMR90 contact maps to achieve *in silico* contact maps with total contacts between 10^4^ and 10^8^. Across the range of total contacts, the maximum entropy approach outperforms the GNN approach based on SCC [Fig. S5C]. For both approaches, the SCC plateaus for read depths greater than 10^6^ but decays relatively rapidly for lower read depths. When using the HiC-Spector metric, the GNN and maximum entropy approaches perform comparably for total contacts between 10^6^ and 10^8^ [Fig. S5D]. For total contacts less than 10^6^, the GNN approach’s performance on the HiC-Spector metric degrades faster than the maximum entropy approach’s performance. Note that the simulated contact maps on which the GNN was trained typically contain 10^6^-10^7^ total contacts. As a result, it is unsurprising that the GNN performance deteriorates below a read depth of 10^6^.

Future work will be needed to extend the GNN to be successful on very sparse Hi-C data, as is common in the single-cell Hi-C setting. Single-cell Hi-C experiments typically yield 10^4^ to 10^6^ total contacts *per cell*. Assuming contacts are distributed uniformly across the genome, the 25.6 Mb regions here would only have 10^2^ to 10^4^ total contacts.

## S4 Relationship to existing machine learning methods

The architecture we use [5] is an example of a graph attention network. Similar to transformer architectures, graph attention networks rely on attention mechanisms [53] as their core building block. Notably, AlphaFold is a transformer architecture that has been remarkably successful in the protein structure prediction domain [1, 20, 24].

Since chromatin and proteins are both biological polymers, one might wonder if we could adapt AlphaFold to predict chromatin structure given appropriate inputs. Our approach has two key advantages over an AlphaFold-esque approach in the chromatin structure prediction setting. First, our approach can obtain accurate structure estimates with far less experimental training data. This is essential for chromatin structure prediction since we don’t have large-scale experimental databases such as those available for protein structure [3]. Second, by using our polymer model, we can map an experimental contact map to a *distribution* of candidate structures rather than a *single* static structure, better reflecting the dynamic nature of the molecules. Effectively, our entire approach can be regarded as a physics-informed generative model for structures instead of a point estimate of a single structure, reflecting both (a) uncertainty in structure estimates and (b) the dynamic nature of chromatin.

## S5 Role of machine learning in molecular simulations

In computational chemistry and biology, coarse-grained particle-based simulation models are essential for studying the structural properties of polymers and large biomolecules. There are two basic frameworks for constructing coarse-grained models, referred to as ‘bottom-up’ and ‘top-down’ [34]. In the bottom-up framework, a coarse-grained model is parametrized to approximate an underlying fine-resolution model. Machine learning is commonly used in the bottom-up framework to learn effective coarse-grained potentials [15, 12].

On the other hand, the top-down framework does not require an underlying fine-resolution model. In the top-down framework, a model is constructed with user-defined energy functional consisting of unknown parameters that must be estimated from experimental data [34]. Our approach falls into this category. In the context of machine learning, top-down simulations can be considered an inverse problem where experimental data is observed, but the parameters of the underlying energy functional are unobserved. There has been relatively little work developing machine learning approaches to parametrize coarse-grained models in the top-down framework [51]. We hope that our approach spurs further development for machine learning tools in the top-down setting.

## S6 Metrics for Comparing Contact Maps

The first metric we consider is the stratum-adjusted correlation coefficient (SCC), which is a weighted average of the Pearson correlation between pairs of off-diagonals of two Hi-C contact maps [55]. To compute the SCC, we wrote our own implementation which operates on dense matrices. The SCC requires two userdefined parameters: a smoothing parameter, *h*, and the maximum range of interactions to consider (i.e., the maximum off-diagonal to consider, *K*). We consider only contacts within the range of 0-5 Mb, as in the original method from Yang et al. [55]. This corresponds to the first *K* = 100 off-diagonals for our resolution of 50 kb. For smoothing, we use a 2D mean filter with a span size of *h* = 5. We choose *h* = 5 because our resolution of 50 kb is closest to the recommendation of using *h* = 5 for 40 kb resolution from Yang et al. We show the effect of varying the filter span size, *h*, (while *K* = 100) in Fig. S7.

As an alternative metric, we consider the Hi-C Spector score [54]. Given two Hi-C contact maps, the Hi-C Spector score first converts each contact map into a graph Laplacian matrix. The Hi-C Spector score is a scaled sum of the Euclidean norm between the first *κ* Laplacian eigenvectors. The metric is scaled to the range [0, 1], where 1 is better. We compute the HiC-Spector score using the implementation provided in https://github.com/gersteinlab/HiC-spector with the first *κ* = 10 Laplacian eigenvectors. Empirically, past work has found the Hi-C Spector score to be more robust to the Hi-C data quality (ie., sequencing read depth) than the SCC [56].

As seen in Table 1 and described in Section 2.2, the maximum entropy approach outperforms the GNN approach based upon the SCC (the Hi-C Spector score is comparable). Despite this, we find that the maximum entropy simulated contact maps often have noticeable visual ‘imperfections’. In Fig. S8, we show four example contact maps where the maximum entropy approach tends to overestimate experimental contact frequencies in some portions of the contact map. The corresponding genomic-distance normalized contact maps are shown in Fig. S9. Note that these examples are not curated; they are the first four contact maps in our experimental test set - all from chromosome 2 of the IMR90 cell line. The significance of these imperfections is unclear, and therefore we include two additional metrics, Corr 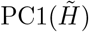 and 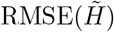

Corr 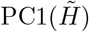 and 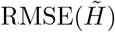 are both measures of the global similarity of genomic-distance normalized contact maps. In contrast, both SCC and HiC-Spector focus primarily on contacts near the main diagonal. By design, the SCC only focuses on contacts within 5 Mb. In addition, the SCC gives higher weight to contacts near the main diagonal. HiC-Spector also focuses on contacts near the main diagonal since it treats the contact map as a graph and compares Laplacian eigenvectors. The primary signal in this graph Laplacian will be the main diagonal of the contact map.

Depending on the application, it may be preferable to focus on the main diagonal (where the majority of contacts are located) or to focus on the compartmentalization pattern. Visually, the GNN approach seems to do better at reproducing long-range compartmentalization patterns [Fig. S8, Fig. S9]. This is supported by the GNN’s better performance on the Corr 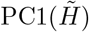 and 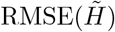 metrics [Table 1].

The origin of this performance difference may be a consequence of differences in the GNN loss function and maximum entropy optimization objective. We train the GNN with mean-squared error loss between the ground truth and estimated 𝒰 matrices. As a consequence, the GNN gives equal weight to all entries in 𝒰. This may explain why the GNN performs best on global measures of contact map similarity. On the other hand, the maximum entropy optimization objective seeks to constrain the average contact frequencies between beads of type *I* and *J* to match between the simulation and experiment (see Section S2.2 for details of the maximum entropy approach). Since the majority of contacts are located near the main diagonal, the maximum entropy approach prioritizes performance near the main diagonal.

## S7 Supplementary Figures

**Table S2:** Neural network ablation results. See Section S2.4.3 for a description of each method. All results in the body of the paper correspond to the baseline method. MSE is the average mean-squared error on the validation set of 1000 simulated contact maps and synthetic parameters. The MSE is reported after log-transformation of 𝒰 to 𝒰^*†*^ as described in Section 4.4. SCC, HiC-Spector, Corr 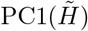, and 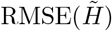 are computed between each experimental contact map and the contact map simulated using the GNN-predicted parameters, averaged over 33 experimental contact maps from the odd chromosomes of the IMR90 cell line.

| Method | MSE | SCC | HiC-Spector | Corr PC1($\tilde{H}$) | RMSE($\tilde{H}$) |
| --- | --- | --- | --- | --- | --- |
| Baseline | 0.406 | 0.725 | 0.533 | 0.913 | 0.677 |
| No log-transform in loss | NA | 0.573 | 0.468 | 0.620 | 0.805 |
| 50 kb resolution | 0.313 | 0.650 | 0.503 | 0.825 | 0.682 |
| Without SignNet (with eigenvectors) | 0.501 | 0.462 | 0.421 | 0.827 | 0.867 |
| Without SignNet (without eigenvectors) | 0.484 | 0.647 | 0.468 | 0.809 | 0.707 |
| Without mean(diagonal($H, \vert i - j\vert $)) in $E_{ij}$ | 0.470 | 0.457 | 0.432 | 0.738 | 0.802 |
| Original message passing layer from [5] | 1.784 | -0.003 | 0.379 | 0.204 | 0.843 |

**Table S3:** Average results for experimental contact maps from even chromosomes of GM12878, HMEC, HUVEC, HAP1, and CH12.LX (GNN was trained only using the IMR90 cell line). Metrics are defined as in Table 1. Maximum entropy optimization was performed with a convergence criterion of *ϵ* = 1*e* − 2. All values are mean *±* standard error.

| Method | SCC | HiC-Spector | Corr PC1($\tilde{H}$) | RMSE($\tilde{H}$) | Simulation Time |
| --- | --- | --- | --- | --- | --- |
| GM12878, HMEC, HUVEC, HAP1 |  |  |  |  |  |
| GNN | $0.71 \pm 0.01$ | $0.54 \pm 0.01$ | $0.85 \pm 0.01$ | $0.70 \pm 0.01$ | $11.4 \pm 0.1$ |
| Maximum Entropy | $0.82 \pm 0.00$ | $0.52 \pm 0.01$ | $0.79 \pm 0.01$ | $1.07 \pm 0.01$ | $75.3 \pm 2.0$ |
| GM12878 |  |  |  |  |  |
| GNN | $0.74 \pm 0.01$ | $0.55 \pm 0.02$ | $0.86 \pm 0.02$ | $0.49 \pm 0.01$ | $9.5 \pm 0.1$ |
| Maximum Entropy | $0.83 \pm 0.01$ | $0.52 \pm 0.02$ | $0.84 \pm 0.02$ | $0.92 \pm 0.03$ | $72.8 \pm 4.1$ |
| HMEC |  |  |  |  |  |
| GNN | $0.66 \pm 0.02$ | $0.53 \pm 0.03$ | $0.82 \pm 0.03$ | $0.7 \pm 0.01$ | $13.5 \pm 0.2$ |
| Maximum Entropy | $0.83 \pm 0.01$ | $0.55 \pm 0.03$ | $0.76 \pm 0.04$ | $0.95 \pm 0.02$ | $77.9 \pm 5.3$ |
| HUVEC |  |  |  |  |  |
| GNN | $0.66 \pm 0.02$ | $0.53 \pm 0.03$ | $0.82 \pm 0.03$ | $0.7 \pm 0.01$ | $13.5 \pm 0.2$ |
| Maximum Entropy | $0.83 \pm 0.01$ | $0.55 \pm 0.03$ | $0.76 \pm 0.04$ | $0.95 \pm 0.02$ | $77.9 \pm 5.3$ |
| HAP1 |  |  |  |  |  |
| GNN | $0.7 \pm 0.01$ | $0.56 \pm 0.02$ | $0.83 \pm 0.03$ | $0.76 \pm 0.01$ | $11.0 \pm 0.3$ |
| Maximum Entropy | $0.82 \pm 0.01$ | $0.54 \pm 0.02$ | $0.79 \pm 0.02$ | $1.18 \pm 0.03$ | $84.3 \pm 4.5$ |
| CH12.LX |  |  |  |  |  |
| GNN | $0.78 \pm 0.01$ | $0.42 \pm 0.02$ | $0.8 \pm 0.04$ | $0.84 \pm 0.02$ | $9.2 \pm 0.2$ |
| Maximum Entropy | $0.81 \pm 0.01$ | $0.38 \pm 0.02$ | $0.85 \pm 0.01$ | $1.29 \pm 0.02$ | $77.0 \pm 4.6$ |

**Table S4:**
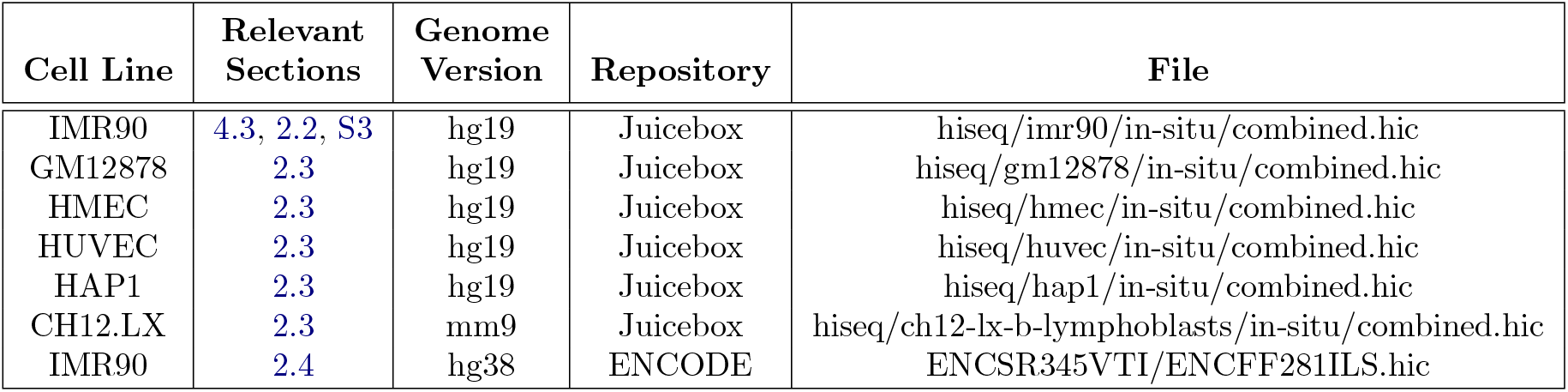
Experimental data sources used in this manuscript. The Juicebox repository can be found at https://www.aidenlab.org/data.html. The ENCODE repository can be found at https://www.encodeproject.org/.

**Figure S1:**
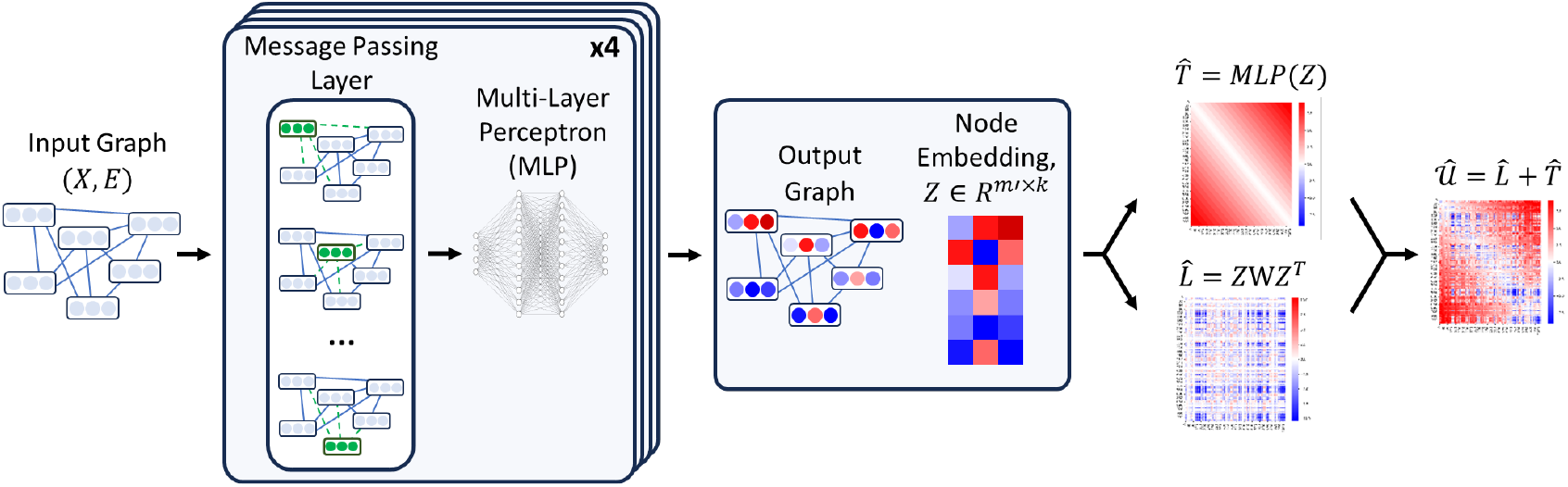
Schematic overview of GNN architecture. See Section 4.4.1 for details. We define a graph (not to scale), node features (*X*), and edge features (*ϵ*) based on a Hi-C contact map. The core graph neural network comprises a message passing layer and a multi-layer perception (MLP). After four iterations, the GNN yields a node embedding, *Z*. The node embedding is used to predict 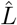 via a bilinear function and 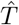 via another MLP. 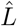 and 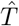 are summed (element-wise) to yield 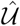.

**Figure S2:**
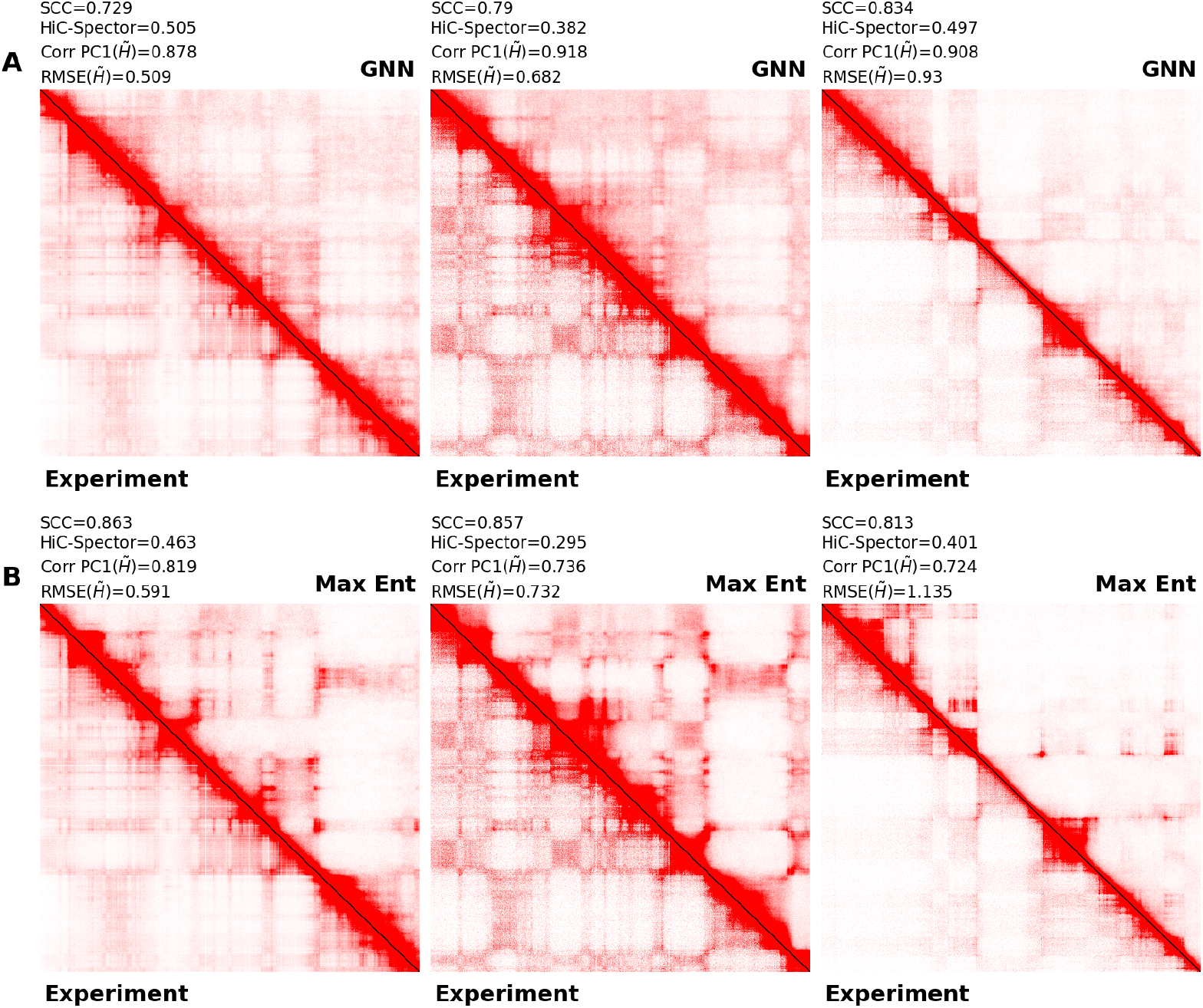
Comparison of GNN and maximum entropy simulated contact maps vs. experimental contact map of Chr2:5.1-30.7 Mb from three cell lines. *Left:* GM12878 human lymphoblastoid cell line. *Middle:* HMEC human mammary epithelial cell line. *Right:* HUVEC human umbilical vein endothelial cell line. The lower triangle shows the experiment. The color bar is the same for all subfigures. **A** The upper triangle shows the GNN simulation. **B** The upper triangle shows the maximum entropy simulation.

**Figure S3:**
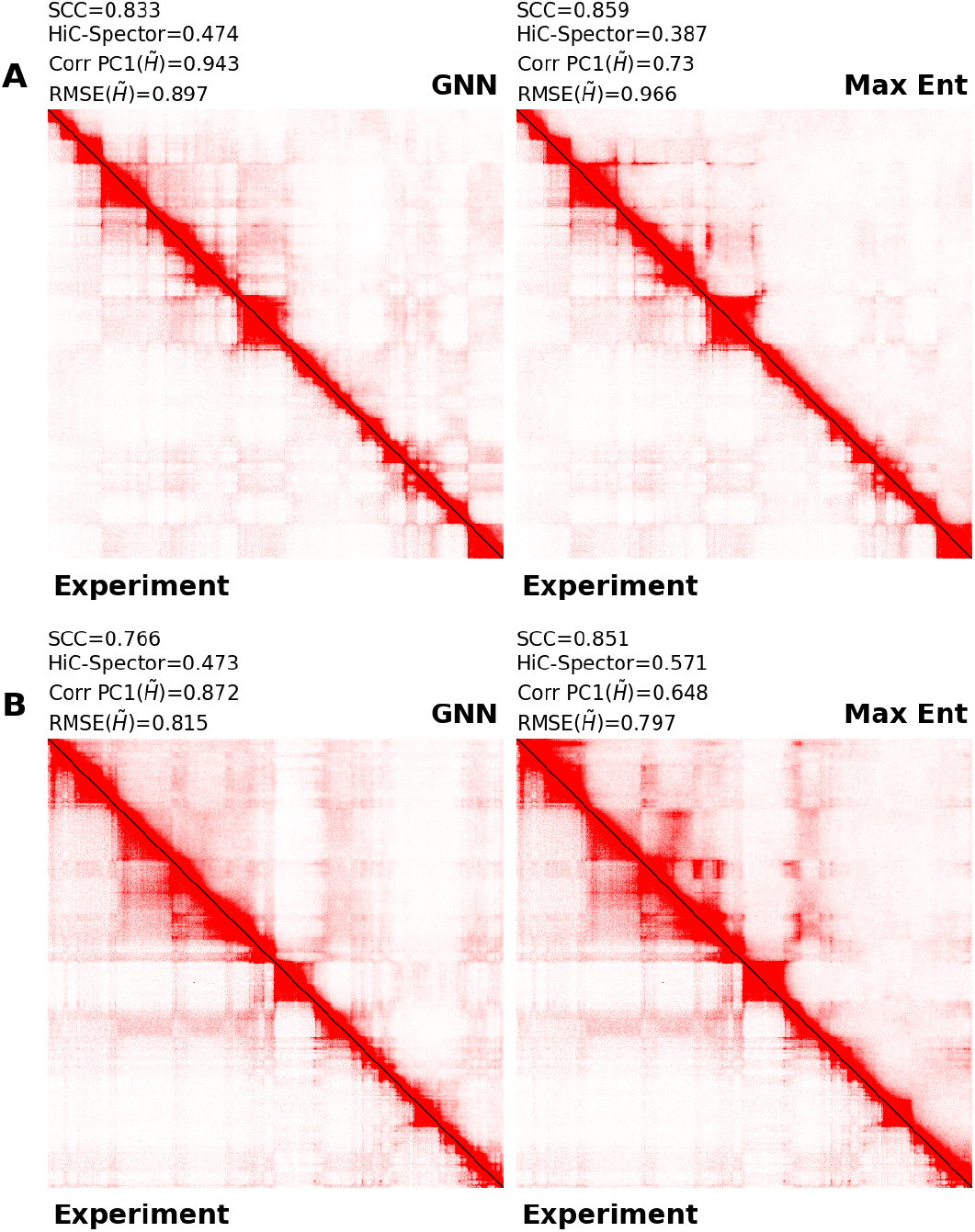
Contact maps corresponding to distances analysis in Fig. 4. **A** IMR90 Chr2:10-25.6 Mb. **B** IMR90 Chr21:14-39.6 Mb. The lower triangle shows the experimental contact map. *Left:* The upper triangle shows the GNN simulated contact map. *Right:* The upper triangle shows the maximum entropy simulated contact map.

**Figure S4:**
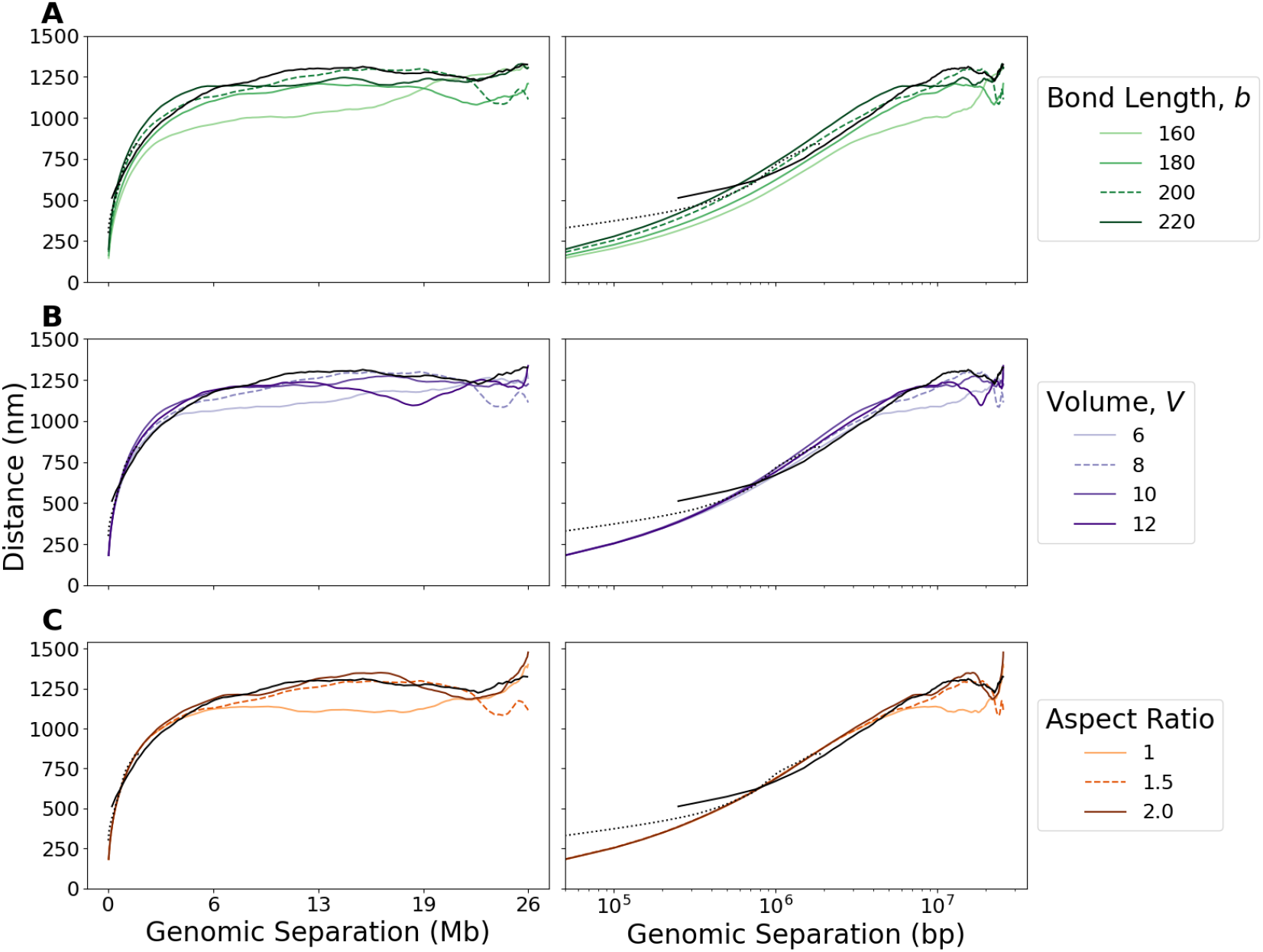
Mean distance scaling as function of genomic separation. The y-axis is the mean euclidean distance averaged over structures and averaged over pairs of loci at a given genomic separation. The two black curves are experimental distance scaling for IMR90 Chr2:10-25.6 Mb. The solid black curve is at 50 kb resolution from [49], and the dotted black curve is at 30 kb resolution from [4]. Colored curves are simulated distance scalings for maximum entropy optimized simulations. The dashed curve in each panel corresponds to the parameters used in the main text. *Left:* Linear scale x-axis *Right:* Log scale x-axis **A** Varying bond length, *b*, holding *V* = 8 *µ*m^3^ and aspect ratio = 1.5 **B** Varying simulation volume, *V*, holding *b* = 200 nm^3^ and aspect ratio = 1.5 **C** Varying aspect ratio holding *b* = 200 nm^3^ and *V* = 8 *µ*m^3^

**Figure S5:**
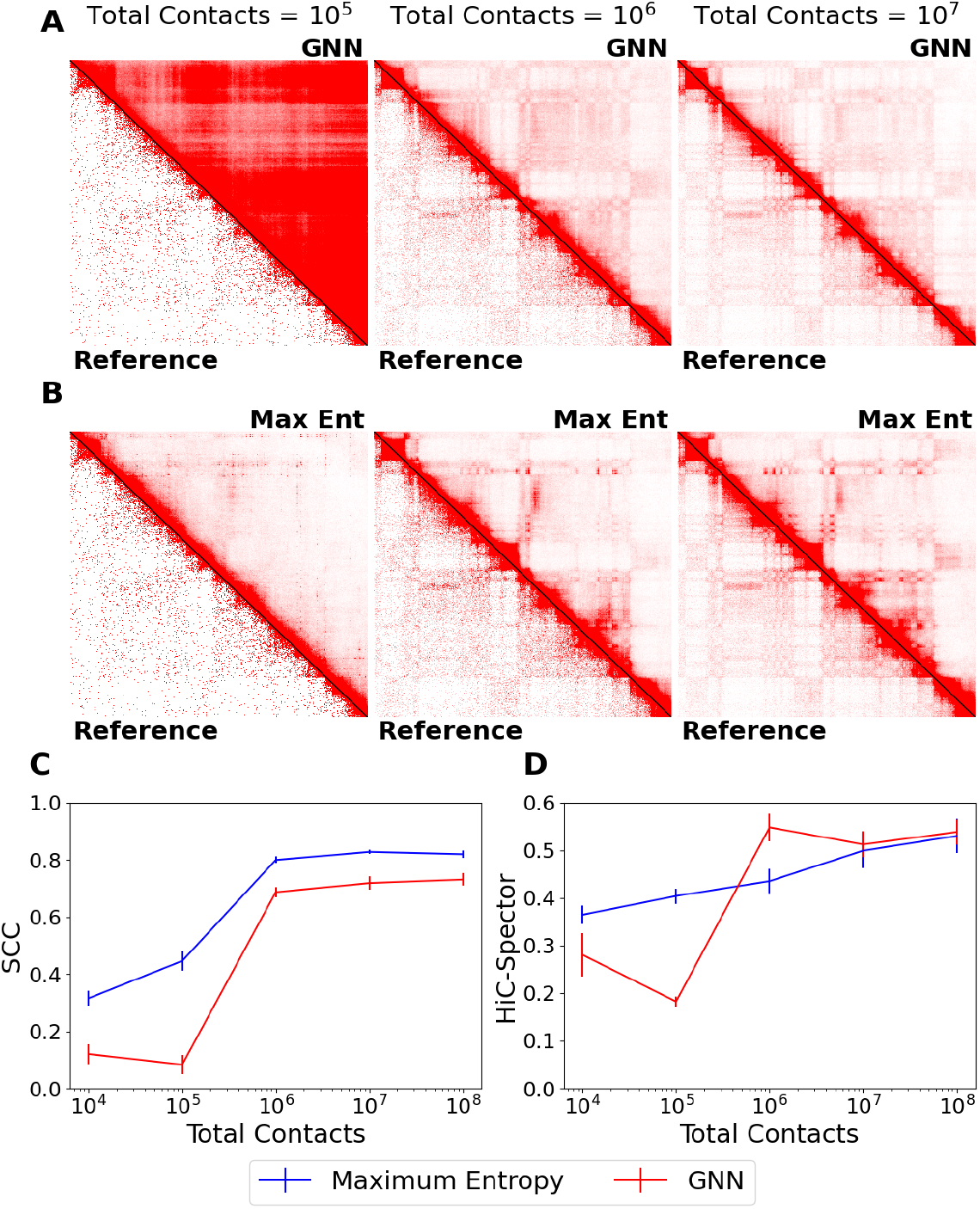
Effect of contact map quality on GNN and maximum entropy performance. **A-B** Contact maps of IMR90 Chr2:149.75-175.35 Mb at different numbers of total contacts. *Left:* total contacts of 10^5^. *Middle:* total contacts of 10^6^. *Right:* total contacts of 10^7^. The lower triangle shows the experimental contact map subsampled to the indicated total contacts. The color bar is the same for all subfigures. **A** The upper triangle is the GNN simulated contact map. **B** The upper triangle is the maximum entropy simulated contact map. **C-D** Average SCC (**C**) and HiC-Spector score (**D**) across ten IMR90 contact maps as a function of total contacts for GNN (red) and maximum entropy (blue) simulations. Error bar shows one ± standard deviation.

**Figure S6:**
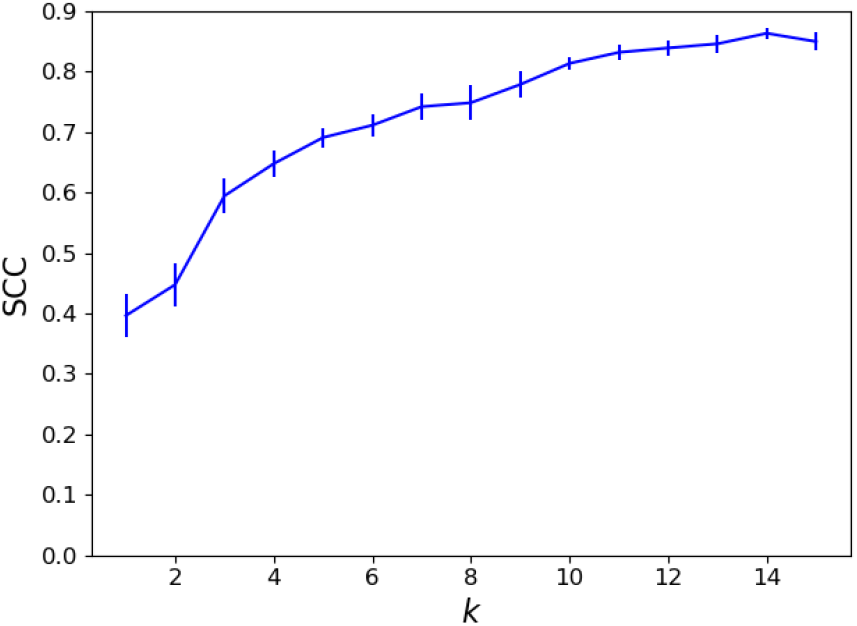
Effect of the number of principal components, *k*, used in the maximum entropy approach on the SCC between experimental and simulated contact maps. SCC is the average SCC from 10 contact maps in the experimental validation set of contact maps from the odd-chromosomes of the IMR90 cell line. Error bars show standard error.

**Figure S7:**
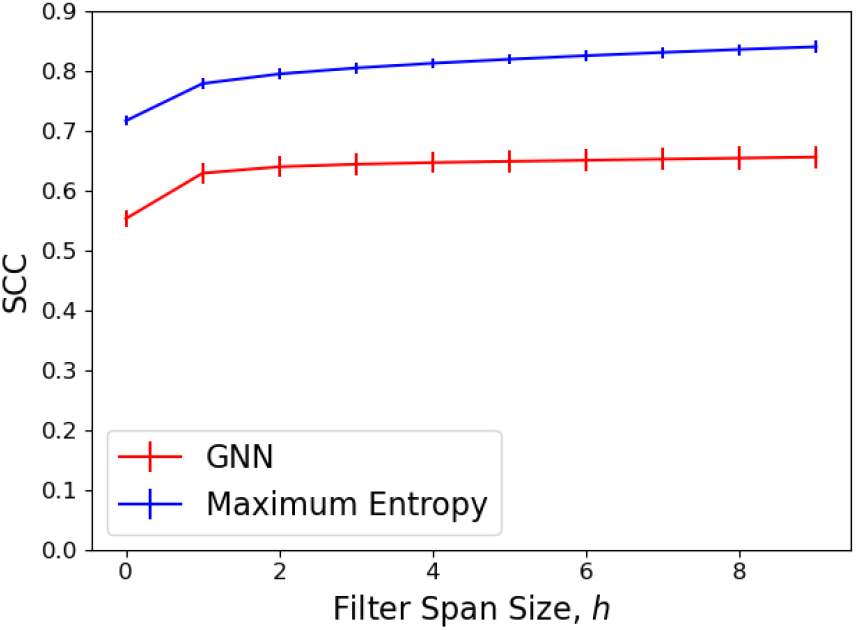
Effect of the filter span size, *h*, when calculating SCC. See Section S6 for details. SCC is the average SCC from 40 contact maps in the experimental test set of contact maps from the even-chromosomes of the IMR90 cell line. SCC is reported for both the GNN approach (red) and maximum entropy approach (blue). Error bars show standard error.

**Figure S8:**
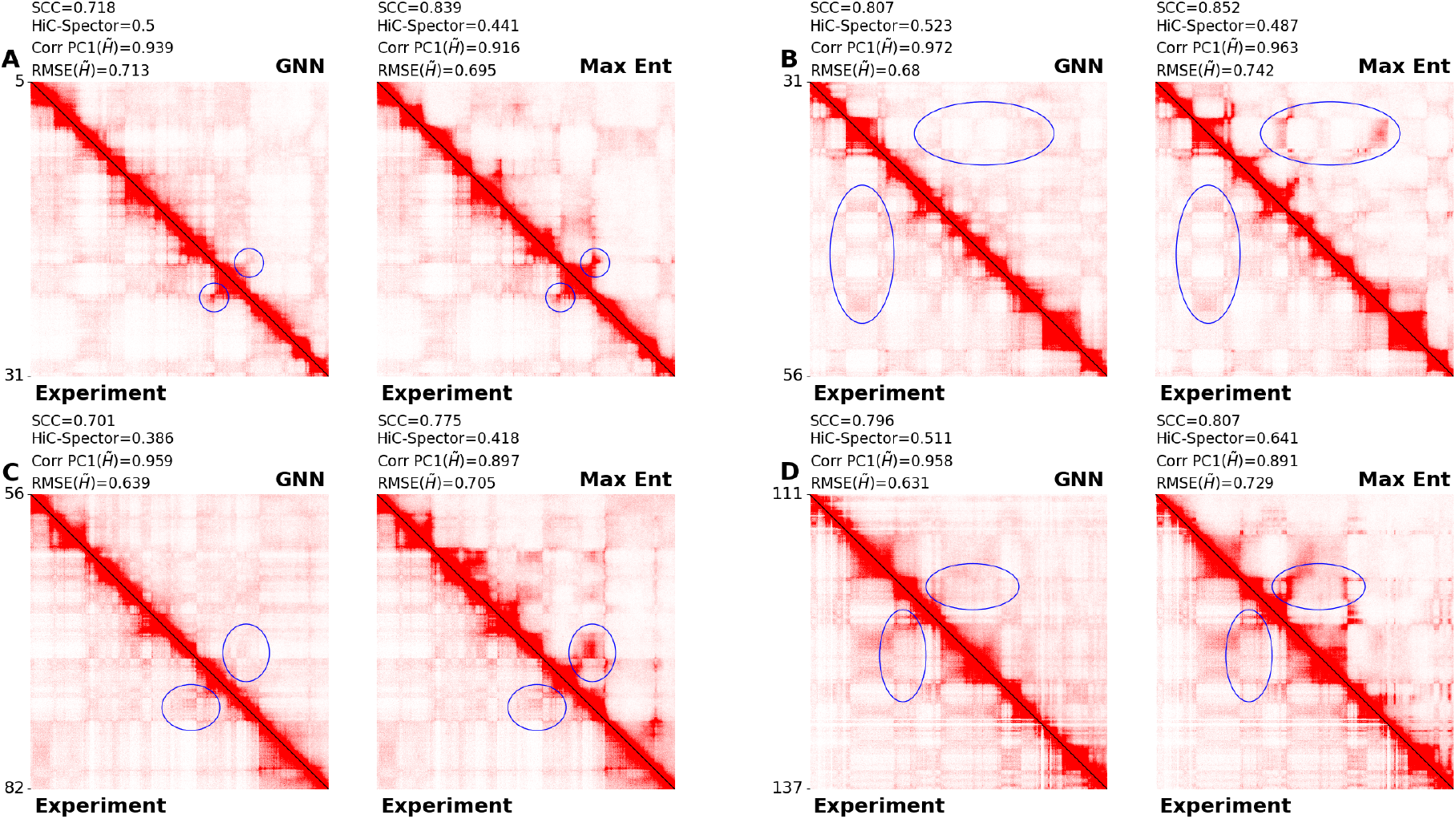
Comparison of GNN and maximum entropy simulated contact maps vs. experimental contact maps. The experimental contact map is always shown in the lower triangle. *Left:* upper triangle is GNN. *Right:* upper triangle is maximum entropy. Blue ellipses highlight regions for which the maximum entropy simulation overestimates the experimental contact frequencies. **A** IMR90 Chr2:5.1-30.7 Mb. **B** IMR90 Chr2:30.7-56.3 Mb. **C** IMR90 Chr2:56.3-81.9 Mb. **D** IMR90 Chr2:111.3-136.9 Mb.

**Figure S9:**
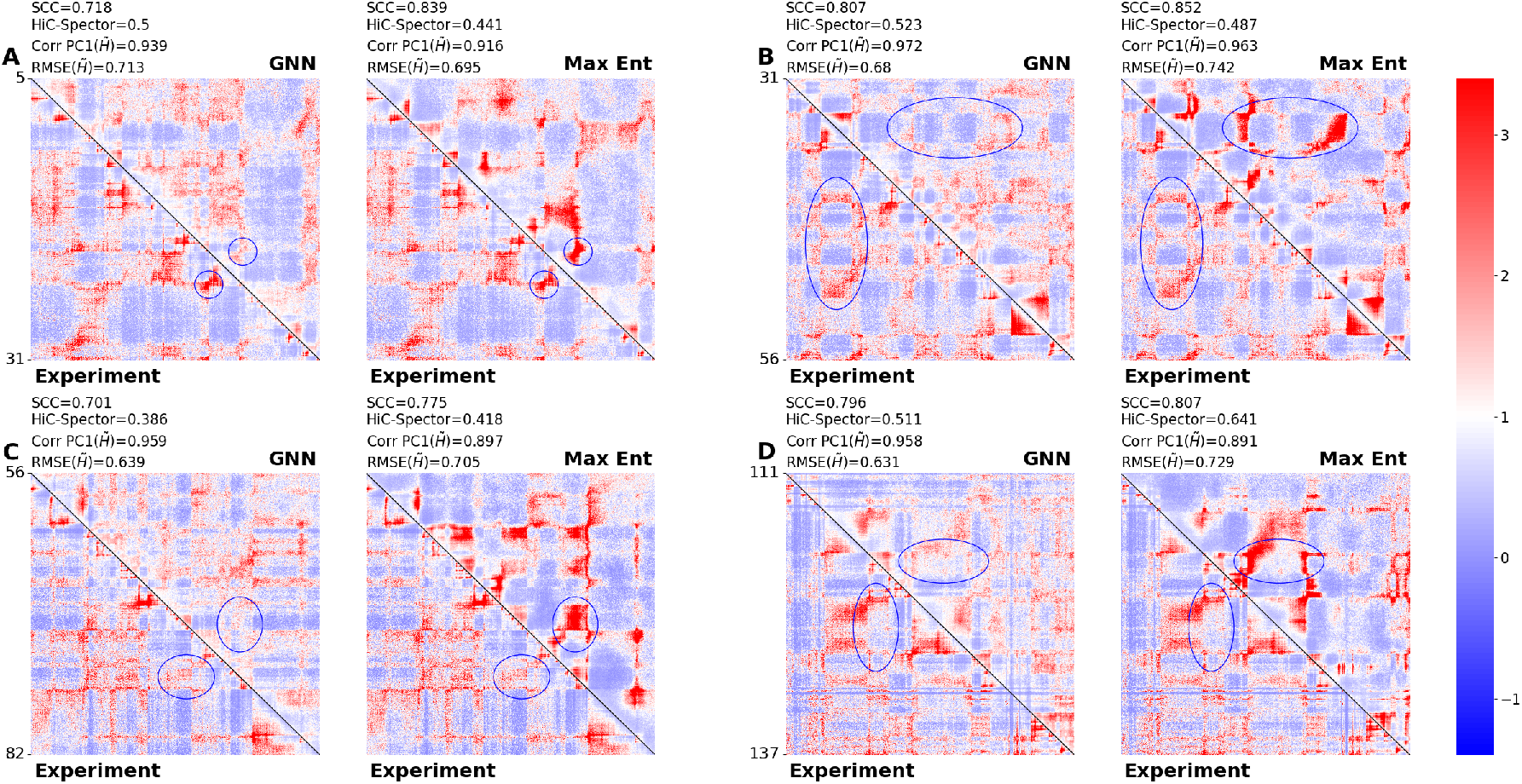
Comparison of genomic-distance normalized contact maps as in Fig. S8. The experimental contact map is always shown in the lower triangle. *Left:* upper triangle is GNN. *Right:* upper triangle is maximum entropy. Blue ellipses highlight regions for which the maximum entropy simulation overestimates the experimental contact frequencies. **A** IMR90 Chr2:5.1-30.7 Mb. **B** IMR90 Chr2:30.7-56.3 Mb. **C** IMR90 Chr2:56.3-81.9 Mb. **D** IMR90 Chr2:111.3-136.9 Mb.

